# Elevated Levels of Mcm7 Disrupt Microtubule and Nucleolar Homeostasis to Drive Genome Instability and Cell Death

**DOI:** 10.64898/2026.07.31.742030

**Authors:** Srijana Dutta, Md. Hashim Reza, Supriya Varsha Bhagat, Tapas Kumar Kundu, Kaustuv Sanyal

## Abstract

Overexpression of pre-replication complex (pre-RC) components is frequently associated with poor prognosis in many types of cancer, yet the mechanisms underlying their pathological effects remain unclear. Using the model yeast *Candida albicans*, we investigated the consequences of elevated Mcm7 levels, a conserved pre-RC subunit previously linked to <u>c</u>hromosome <u>in</u>stability (CIN). Unlike other pre-RC components, Mcm7 overexpression severely compromises nuclear integrity, resulting in chromatin fragmentation, DNA double-strand breaks, and cell death. Excess Mcm7 disrupts chromosome segregation by impairing microtubule function, disrupting centromere organization, generating abnormal spindle structures, and causing defective nuclear migration, collectively leading to mitotic failure. We further discovered ectopic accumulation of Mcm7 within the nucleolus, accompanied by widespread transcriptional disruption of nucleolus-associated genes. Supporting the relevance of these findings to human disease, cancer transcriptomic datasets reveal a strong association between *MCM7* expression and nucleolar gene expression programs, while human cancer cell lines with higher MCM7 expression exhibit increased nucleolar number and size. Together, our findings identify Mcm7 as a previously unrecognized regulator of cytoskeletal and nucleolar homeostasis and uncover an unexpected mechanism by which replication factor dysregulation promotes CIN.

## Introduction

Preservation of genomic stability during cell division is a cardinal requirement for the faithful propagation of life. Errors in DNA replication, DNA repair, or chromosome segregation can give rise to CIN, which is a major factor underlying adverse outcomes such as genetic and developmental diseases and cancer (Vassilev and DePamphilis 2017; Hosea et al. 2024). Hence, it is imperative to delineate the guardians of genome stability; factors whose dysregulation is not tolerated by cells. A recent study in the diploid budding yeast, *Candida albicans,* screened 1067 genes and reported 6 genes whose overexpression led to loss <u>o</u>f <u>h</u>eterozygosity (LOH), an event driven by CIN (Jaitly et al. 2022). The replicative helicase subunit Mcm7 was among the 6 genes, and its overexpression led to cell cycle arrest; however, the basis of this dosage intolerance has not been characterized.

Mcm7 is a member of the 6-subunit <u>m</u>ini<u>c</u>hromosome <u>m</u>aintenance <u>c</u>omplex (MCM2-7), the DNA helicase machinery required for DNA replication. MCM, in turn, is part of the pre-RC, which consists of the <u>o</u>rigin recognition complex (ORC1-6), Cdc6, and Cdt1 (the MCM loader) (Fig. 1A). The pre-RC is responsible for licensing replication origins for firing and ensuring that DNA replication occurs once and only once per cell cycle (Nishitani and Lygerou 2002). This regulation is critical, as both under-replication and re-replication induce genome instability (Nishitani and Lygerou 2002; DePamphilis 2006; Sclafani and Holzen 2007). Besides DNA replication, the involvement of pre-RC members in chromosome segregation is also emerging. For instance, Mcm2 acts as a co-chaperone for the centromere-specific histone H3 variant CENPA, along with its principal chaperone, HJURP (Zasadzińska et al. 2018). Cdt1 in humans associates with Ndc80, an outer kinetochore protein, mediating stable kinetochore-microtubule attachment, which is required for timely and precise chromosome segregation (Varma et al. 2012; Agarwal et al. 2018). Mcm2 and Orc4 are essential for centromere stabilization in *C. albicans*, as their loss leads to the degradation of CENPA^Cse4^ (Sreekumar et al. 2021). These observations suggest that dysregulation of pre-RC proteins may influence genome stability through pathways extending beyond replication initiation.

**Figure 1.**
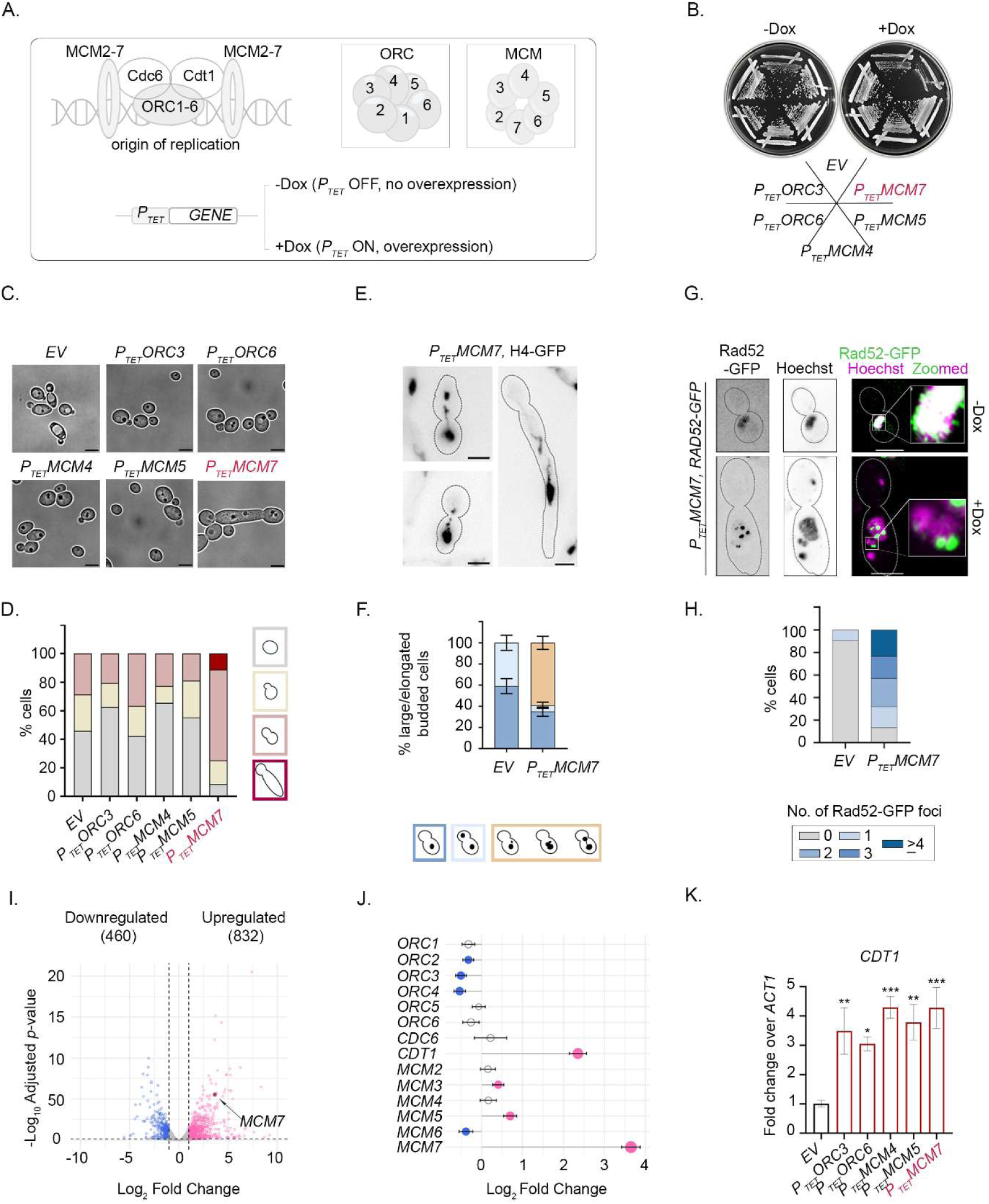
Chromosomal instability upon Mcm7 overexpression results in cell death. A) Schematic portraying the pre-RC components and the conditional overexpression strategy employed in the study. The gene of interest is placed downstream of the inducible tetracycline promoter (*P_TET_*) in an ectopic safe haven locus (*RP10*). The addition of doxycycline (Dox) induces overexpression from *P_TET_*. B) Overexpression-mediated cell lethality evaluated for individual pre-RC components (*EV*- CaPJ160 *P_TET_ORC3*- SRICa007; *P_TET_ORC6*- SRICa012; *P_TET_MCM4*- SRICa016; *P_TET_MCM5*- SRICa020; *P_TET_MCM7*- CaPJ165) by streaking strains on YPDU plates with either no Dox (-Dox) or addition of Dox (+Dox), wherein Dox induces overexpression. Plates were imaged after 48 h of incubation at 30 °C. C) Brightfield microscopy images showing the morphological differences of *C. albicans* cells upon overexpression of individual pre-RC subunits. Images captured after treating cells with Dox for 8 h. Scale bar: 5 µm. D) Quantification of the distribution of different cell cycle stages upon overexpression of pre-RC subunits. *n* ≥ 100 cells. E) Fluorescence microscopy images of H4-GFP upon Mcm7 overexpression (SRICa178), representing the abnormal nuclear morphology observed. Scale bar: 5 µm. F) Quantification of nuclear distribution in large-budded and elongated-budded cells in *EV* (SRICa175) and *P_TET_MCM7* (SRICa178) strains, treated with Dox for 8 h. *N* = 3, *n* ≥ 100 for each biological replicate. G) Fluorescence microscopy images showing Rad52-GFP localization with respect to the nucleus stained with Hoechst in *P_TET_MCM7* cells (SRICa234) with or without the addition of Dox. Scale bar: 5 µm. H) Quantification of the number of Rad52-GFP foci in *EV* (SRICa265) and *P_TET_MCM7* (SRICa234), both treated with Dox for 8 h. *n* ≥ 100 cells. I) Volcano plot depicting the number of genes differentially expressed in *P_TET_MCM7* cells (CaPJ165) compared to *EV* control (CaPJ160). Analysis conditions: log2 fold change cutoff at -1 and 1, *p*-adjusted value cutoff at 0.05. *MCM7* is highlighted as a maroon circle to mark its position in the upregulated wing of the volcano plot. J) Lollipop graph generated from RNA-seq experiment, depicting fold-change in the expression of all pre-RC subunits in *P_TET_MCM7* cells compared to *EV* control. Lollipops at the positive *x*-axis or with pink heads are up-regulated, whereas lollipops with blue heads or towards the negative *x*-axis are down-regulated. Non-coloured lollipop heads are non-significant. K) Real-time PCR quantification of *CDT1* transcripts in *EV* and respective pre-RC overexpression strains, after treatment with Dox. Fold change relative to *ACT1* has been plotted. *N* = 3, Ordinary one-way ANOVA, multiple comparison with control group *EV*, *P*-value: (***) <0.001, (**) < 0.01, (*) < 0.05. Error bars indicate standard <u>e</u>rror of <u>m</u>ean (SEM).

Loss-of-function studies have firmly established the essential requirement for pre-RC proteins during DNA replication, whereas considerably less is known about the consequences of increasing their abundance. Consistent with their essential role in proliferation, elevated expression of pre-RC components is frequently observed in highly proliferative cancers and is widely used as a diagnostic and prognostic marker (Wang and Liu 2023; Zhao et al. 2024). While overexpression may be a collateral consequence of extensive genomic instability during cancer progression, reports also suggest that dysregulation of individual pre-RC subunits can actively contribute to genomic instability and tumorigenesis (Lau et al. 2007; Yu et al. 2020; Lattmann et al. 2022). It was reported that increased MCM7 expression is associated with relapses, local invasion, and severe tumor grade in prostate cancer. Furthermore, constitutive Mcm7 expression was sufficient to drive increased DNA synthesis, cell proliferation, and cell invasion capacity (Ren et al. 2006). Elevated MCM7 contributed to tumor formation, progression and malignant conversion in a mouse model, thereby directly implicating MCM7 deregulation in tumorigenesis (Honeycutt et al. 2006). Owing to the multifaceted roles of pre-RC components and their widespread association with cancer, understanding the pleiotropic effects of their overexpression is important.

To address how dysregulation of a replication licensing factor influences genome stability, we investigated the consequences of Mcm7 overexpression in the genetically tractable yeast *C. albicans*, whose highly plastic genome and tolerance to aneuploidy make it an excellent model for studying CIN (Legrand et al. 2019). We demonstrate that, unlike other pre-RC components, elevated Mcm7 profoundly compromises chromosome segregation and cell viability. Mechanistically, our findings implicate microtubule organization and nucleolar homeostasis as key cellular processes perturbed by elevated Mcm7, revealing a broader cellular influence of this conserved replication factor.

## Results

### Increased Mcm7 levels cause cell lethality

While Mcm7 is a member of the pre-RC, which cooperatively functions within a complex to facilitate DNA replication, it is not unusual for pre-RC components to have subunit-specific roles (Forsburg 2004). Thus, having shown a cell cycle arrest phenotype upon Mcm7 overexpression in *C. albicans* (CaPJ165) (Jaitly et al. 2022), we sought to examine whether this is a common phenomenon upon overexpression of any pre-RC subunit or is specific to Mcm7. We generated *C. albicans* strains overexpressing Orc3 (SRICa007) or Orc6 (SRICa012) from the ORC and Mcm4 (SRICa016) or Mcm5 (SRICa020) from the MCM complex, using the inducible tetracycline/doxycycline overexpression plasmid collection (Chauvel et al. 2012) (Fig. 1A). Overexpression of each of the gene transcripts compared to the empty vector control (*EV*, CaPJ160) for respective *C. albicans* transformants was confirmed by real-time PCR. A significantly high expression was observed for each subunit after 8 h of induction with doxycycline (Dox), wherein the addition of Dox induces expression of the genes placed downstream of the tetracycline promoter (Fig. S1A). Strikingly, only excess Mcm7 could induce cell death, while overexpression of other pre-RC subunits did not lead to any visible growth defects (Fig. 1B). Polarised/elongated growth in *C. albicans* has been linked to responses to genotoxic stress and to improper cell cycle progression (Bachewich et al. 2005; Sahoo et al. 2025). Such a morphological transition was observed in cells overexpressing only Mcm7, but not other pre-RC subunits (Fig. 1C). The accumulation of large-budded and elongated-budded stages upon Mcm7 overexpression confirms a cell cycle arrest. On the other hand, the distribution of the cell population was comparable to that of the *EV* control for other pre-RC subunits (Fig. 1D). Therefore, overexpression of Mcm7 uniquely elicits an irreparable cell cycle arrest, an event not induced by other pre-RC components tested.

### Excess Mcm7 causes chromatin damage and transcriptional rewiring

MCMs are DNA-binding proteins that alter the dynamic chromatin structure during DNA replication. To visualize the state of chromatin in the presence of excess Mcm7, we tagged histone H4 with GFP in *EV* control (SRICa175) and *P_TET_MCM7* cells (SRICa178). Incidence of multiple nuclear fragments, stretched/distorted nuclear shape and fine chromatin threads were visible upon Mcm7 overexpression, indicative of genomic catastrophe (Fig. 1E). This class of abnormal nucleus was present in ∼60% of the large/elongated-budded cells as opposed to none observed in *EV* control, accompanied by a decrease in the percentage of large-budded cells with a segregated nucleus (∼6% in *P_TET_MCM7* vs ∼40% in *EV* cells) (Fig. 1F). To determine whether these structural abnormalities reflected underlying DNA damage, we functionally expressed GFP-tagged DNA repair protein Rad52 (SRICa234) and monitored foci formation as a readout of <u>h</u>omologous recombination (HR)-mediated repair (Lisby et al. 2001, 2003). A pan-nuclear signal was observed in the absence of Dox, whereas discrete puncta were observed when Dox was added to induce Mcm7 overexpression (Fig. 1G), suggesting DNA breaks. A large proportion of *P_TET_MCM7* cells harbored more than one Rad52-GFP puncta, clearly indicating widespread DNA damage (Fig. 1H).

There still lies a possibility that overexpression of Mcm7 affects the expression levels of other pre-RCs, since Mcm7 has been reported to act as a co-factor of the transcription factor Mcm1 to modulate its own expression as well as the expression of Cdc6 and Mcm5 in *Saccharomyces cerevisiae* (Fitch et al. 2003). Thus, Mcm7 might induce chromatin damage by modulating the expression of its partners, thereby enhancing the phenotype. RNA-seq was performed on *EV* and *P_TET_MCM7* cells treated with Dox to identify transcriptomic changes upon Mcm7 overexpression. <u>P</u>rincipal <u>c</u>omponent <u>a</u>nalysis (PCA) revealed distinct clustering between *EV* control and *P_TET_MCM7* strain, with minimal differences in PC1 between their biological replicates (Fig. S1B). This suggests that cells have a significantly altered transcriptional landscape when Mcm7 is overexpressed. Compared to housekeeping genes, such as actin (*ACT1*), an expected increase in *MCM7* transcripts was observed in the genome browser view (Fig. S1C), a difference that was statistically significant when normalized transcript counts for *ACT1* and *MCM7* were plotted in both conditions (Fig. S1D). With a log2 fold change cut-off at 1 and -1, and an adjusted *p*-value cut-off at 0.05, 832 genes were found to be upregulated, whereas 460 genes were downregulated in the Mcm7 overexpression condition (Fig. 1I), accounting for ∼21% of the total transcriptome showing altered gene expression. Although the observed changes in the transcriptional profile may be partially attributable to cellular stress associated with cell cycle arrest and Mcm7-induced cell death, the possibility that elevated Mcm7 levels directly elicit transcriptional alterations detrimental to cellular homeostasis cannot be excluded.

Subsequently, we checked the expression profile of other pre-RC subunits upon Mcm7 overexpression. The relative abundance of other pre-RC components remained largely unchanged, with the notable exception of *CDT1*, the MCM helicase loader, which was upregulated (Fig. 1J). We wondered if the concomitant increase of the loader is specific to Mcm7 overexpression, which facilitates its excess recruitment to the genome, resulting in the phenotype. To test this, we examined *CDT1* transcript levels upon overexpression of individual pre-RC subunits. Overexpression of Orc3, Orc6, Mcm4, and Mcm5 also led to elevated *CDT1* transcripts (Fig. 1K). Despite this shared transcriptional response, only excess Mcm7 elicited pronounced cell cycle defects and loss of viability. These observations confirm that the phenotypes associated with Mcm7 overexpression result from perturbation of a subunit-specific regulatory network that is sensitive to Mcm7 dosage.

### Overexpression of Mcm7 activates Mad2-dependent spindle assembly checkpoint

A previous FACS analysis showed >2N DNA content, with a prominent 4N population in *C. albicans* cells overexpressing Mcm7 (Jaitly et al. 2022), indicating that DNA replication was largely completed but the cell cycle was arrested at G2/M. The <u>s</u>pindle <u>a</u>ssembly <u>c</u>heckpoint (SAC) monitors kinetochore-microtubule attachments and restrains anaphase onset until proper bipolar attachment of all chromosomes is achieved. To determine whether cell cycle arrest is mediated by the SAC, we overexpressed Mcm7 in a strain in which a key SAC component, *MAD2*, is deleted (SRICa131). Deletion of *MAD2* renders the SAC inactive. Indeed, we observed a bypass of cell cycle arrest upon SAC inactivation, as reflected by improved growth of cells in SRICa131 compared to cells overexpressing Mcm7 with intact *MAD2* (SRICa064) (Fig. 2A). Such cells proceeded through the cell cycle and eventually lost viability due to erroneous chromosome segregation. As revealed by FACS analysis, the accumulation of 4N DNA content in SRICa064 was partly reverted when SAC was inactivated (SRICa131) (Fig. S2A), confirming that a SAC-mediated arrest contributes to G2/M cell accumulation. Progression through the cell cycle in the absence of the SAC was further established by the decrease in the percentage of cells accumulated in the large/elongated-bud stage, as compared to cells overexpressing Mcm7 with the SAC on (Fig. 2B), with a simultaneous decrease in the percentage of abnormal nuclear morphotypes and an increase in segregated nuclear population (Fig. 2C). The SAC remains active until defects in kinetochore-microtubule attachments arising from altered kinetochore stability or microtubule dynamics are resolved (Fig. S2B). To identify the basis of SAC activation following Mcm7 overexpression, we systematically examined each layer of the chromosome segregation machinery, beginning with the kinetochore, the chromatin-associated structure that mediates microtubule attachment.

**Figure 2.**
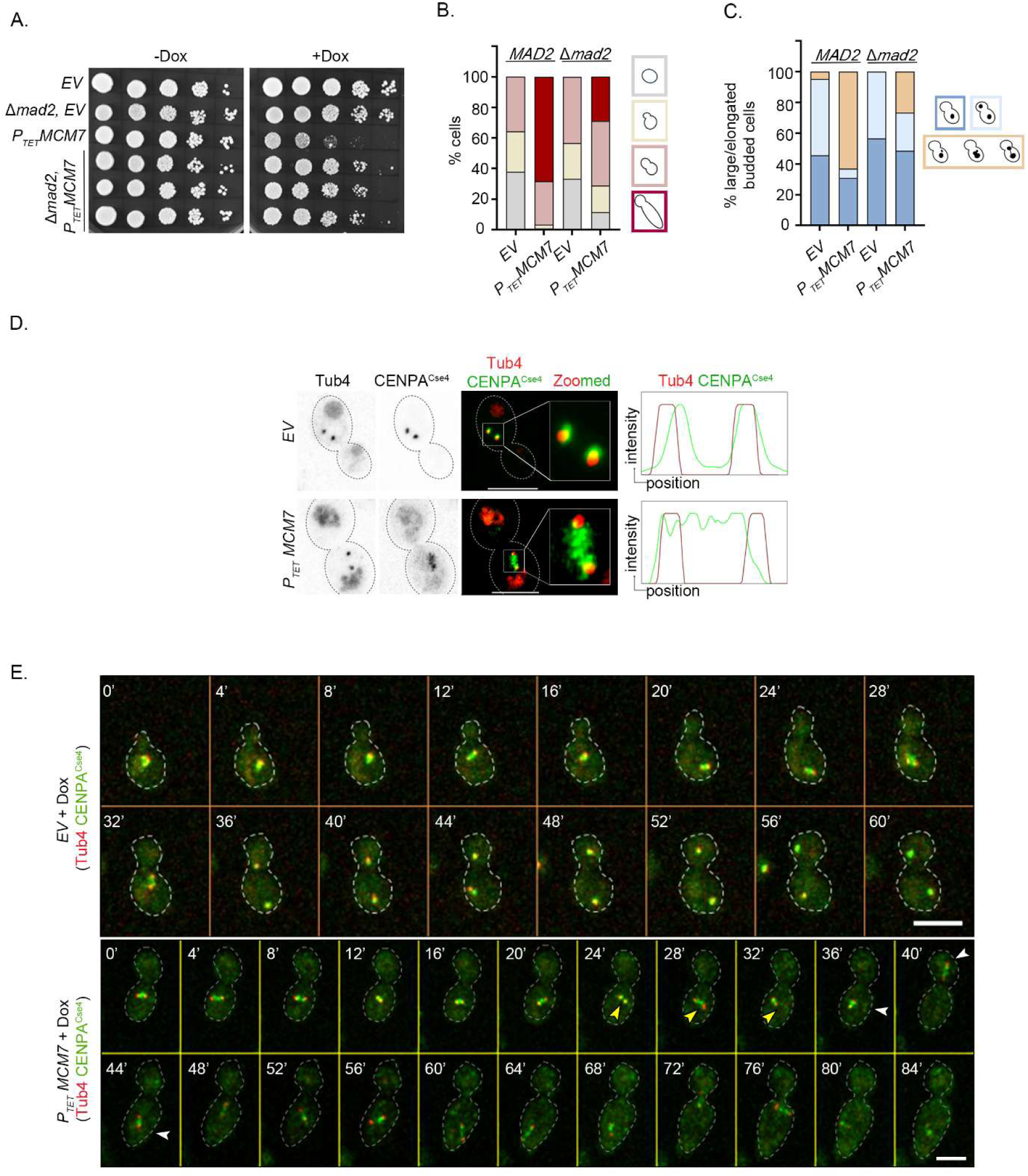
Chromatin instability upon Mcm7 overexpression is accompanied by centromere unclustering and SAC-dependent arrest. A) Ten-fold serial dilution spotting assay to compare growth differences in *P_TET_MCM7* with *MAD2* intact (SRICa064) and Δ*mad2*, *P_TET_MCM7* (SRICa131). *EV* with intact *MAD2* (CaPJ170) and in Δ*mad2* strain (SRICa246) are used as controls. Plates were imaged after 48 h of incubation at 30 °C. B - C) Quantification for the distribution of different cell cycle stages and nuclear segregation pattern, respectively, in *EV* control and upon overexpression of Mcm7 with or without active SAC. *n* ≥ 100 cells. D) Fluorescence microscopy images of CENPA^Cse4^-GFP (centromere marker) and Tub4-mCherry (SPB marker) in *EV* (CaPJ160) and *P_TET_MCM7* (CaPJ165) with comparable spindle length (in *EV*: 1.199 µm and in *P_TET_MCM7*: 1.663 µm), treated with Dox for 8 h. Scale bar: 5 µm. E) Live-cell microscopy montages of Tub4-mCherry and CENPA^Cse4^-GFP, post 8 h of Dox treatment. The top corner represents time in min. Yellow arrowheads: dynamic unclustering of centromeres. White arrowheads: oscillation of the SPB-kinetochore pair between the daughter and mother bud. Scale bar: 5 µm.

### Mcm7 dosage is critical for kinetochore stability and clustering

Kinetochores are generally clustered at most stages of the cell cycle in budding yeasts, and maintenance of the centromere clustering state is crucial for higher-order genome organization (Guin et al. 2020; Dutta et al. 2025; Polisetty et al. 2025). Fluorescence microscopy of centromeric histone CENPA^Cse4^-GFP, marking the kinetochore, and Tub4-mCherry, marking the <u>s</u>pindle <u>p</u>ole <u>b</u>ody (SPB) revealed a stretched, unclustered centromere arrangement, rather than a tight centromere cluster, lodged between duplicated SPB puncta in cells overexpressing Mcm7 (Fig. 2D). This is indicative of an altered chromosomal organization in general and improper centromere clustering in particular. Disruption of kinetochore stability is easily assessed in *C. albicans* by measuring the protein stability of CENPA^Cse4^, which is prone to degradation during kinetochore collapse, often preceded by kinetochore unclustering (Thakur and Sanyal 2012). However, western blotting analysis showed that the cellular levels of CENPA^Cse4^ were unperturbed upon Mcm7 overexpression (Fig. S2C). On the other hand, the levels of the canonical histone H3 were drastically reduced (Fig. S2C). Having observed a drop in the canonical H3 pool and unusual CENPA^Cse4^ signals upon Mcm7 overexpression, we were curious to examine whether CENPA^Cse4^ could spread into neighbouring regions. We analyzed the occupancy pattern of CENPA^Cse4^-TAP at centromere 7 (*CEN7*) by ChIP at 1-kb intervals across the centromeric and pericentric regions and found no statistically significant difference relative to *EV* control in CENPA^Cse4^occupancy, suggesting that kinetochore integrity was preserved upon Mcm7 overexpression (Fig. S2D).

A close functional interaction between the MCM subunit Mcm2 and the kinetochore was reported in *C. albicans*, wherein the absence of Mcm2 led to CENPA^Cse4^ degradation, indicating a role for MCM subunits in maintaining kinetochore integrity (Sreekumar et al. 2021). To test if Mcm7 has a conserved kinetochore function like its sister subunit, Mcm2, we generated a Mcm7 conditional promoter shut-down mutant (SRICa041 and SRICa045) by deleting the first allele and placing the second allele under a regulatable *MET3* promoter (*P_MET3_*), which is repressed by methionine (+M) and cysteine (+C) in the media (Care et al. 1999; Reuss et al. 2004) (Fig. S3A). Mcm7 is critical for cell viability, as is evidenced by the lack of growth in the +M +C plate (non-permissive, NP) (Fig. S3B). Cell cycle analysis showed that cells are accumulated at the large-budded and elongated-budded stages, consistent with its overexpression defects (Fig. S3C). Most large/elongated-budded cells had unsegregated nuclei (∼80%), indicating failure to transit to anaphase (Fig. S3D); however, we did not observe any abnormal nuclear phenotype characteristic of Mcm7 overexpression. Based on the detection limit of the microscope, we observed a complete loss of CENPA^Cse4^-GFP signals in Mcm7-depleted cells (Fig. S3E). Consistent with the findings for Mcm2, the absence of Mcm7 led to reduced CENPA^Cse4^ occupancy at centromeres (Fig. S3F) and CENPA^Cse4^ degradation, as observed by western blotting (Fig. S3G). Hence, while any alteration (overexpression and depletion) in Mcm7 levels result in cell cycle arrest and cell death, depleted protein levels compromise centromere stability, whereas elevated Mcm7 levels impair centromere clustering, suggesting a distinct mechanism of action for the dosage of Mcm7 in maintaining chromosome segregation fidelity.

Next, we investigated the dynamic nature of the clustering defect in Mcm7 overexpression by performing live-cell microscopy on the Tub4-mCherry, CENPA^Cse4^-GFP strain (CaPJ165), and observed two atypical events: a) migration of the duplicated SPBs to the daughter cell and backtracking (∼46.35% cells, 13 out of 28 live-cell movies), and b) dynamic CENPA^Cse4^ stretching (∼39.26% cells, 11 out of 28 live-cell movies) (Fig. 2E, white arrowhead: type a, yellow arrowhead: type b). Mutants of cytoskeletal proteins (*kip3*, *kip1*, and *cin8*) display a similar unclustered kinetochore occupying the spindle equator in *S. cerevisiae* (Tytell and Sorger 2006; Gardner et al. 2008; Wargacki et al. 2010). Thus, we subsequently examined the changes in microtubule organization when Mcm7 is present in excess in cells.

### Mcm7 overexpression remodels the microtubule landscape and perturbs nuclear movement

Microtubules (MTs) in *C. albicans* can be categorized into three classes: a) spindle or nuclear MTs (nMTs), that emanate from the SPBs towards the nuclear interface, b) astral MTs (aMTs) that emanate from SPBs towards the cytoplasmic interface, and c) free cytoplasmic MTs (cMTs), whose origin remains unknown (Barton and Gull 1988). MTs were visualized using a Tub2-GFP (β-tubulin) construct in cells having either *EV* (CaRG080) or *P_TET_MCM7* (SRICa157). Strikingly, an extensive network of aMTs and cMTs was visible in cells overexpressing Mcm7 (Fig. 3A). An increase in the length of the spindle (Fig. 3B), aMTs (Fig. 3C) and cMTs (Fig. 3D) was observed upon quantification, suggesting that excess Mcm7 resulted in lengthening of MTs. Cells subjected to replication poisons such as methyl methanesulfonate (MMS) and hydroxyurea (HU) did not display the aberrant MT properties as observed in the Mcm7 overexpression condition (Fig. S4A), confirming that general replication defects, including induction of double-strand breaks (with the addition of MMS), do not result in altered MT properties. In addition to increased MT length, we observed temporal dysregulation of cMTs during Mcm7 overexpression. *EV* cells treated with Dox displayed a cell cycle stage-specific presence of cMTs, wherein a clear absence of cMTs in the short spindle stage (<1.5µm) until anaphase is apparent, similar to wild-type (Fig. S4B) (Finley and Berman 2005; Reza et al. 2024). In contrast, a pronounced and persistent cMT network is evident upon Mcm7 overexpression despite a metaphase-like spindle length, and most cells fail to enter the anaphase spindle stage (Fig. S4B). Furthermore, increased spindle length necessitates robust maintenance of spindle integrity; failure to do so results in spindle bending and breakage, which are often catastrophic. Cells overexpressing Mcm7 displayed spindle buckling, in which the spindle dynamically bends, deforms, and breaks (Fig. S4C, pink arrowheads), while cells with *EV* did not show any MT buckling behavior during cell cycle progression (Fig. S4C). Collectively, a disruption of MT homeostasis is evident upon Mcm7 overexpression.

**Figure 3.**
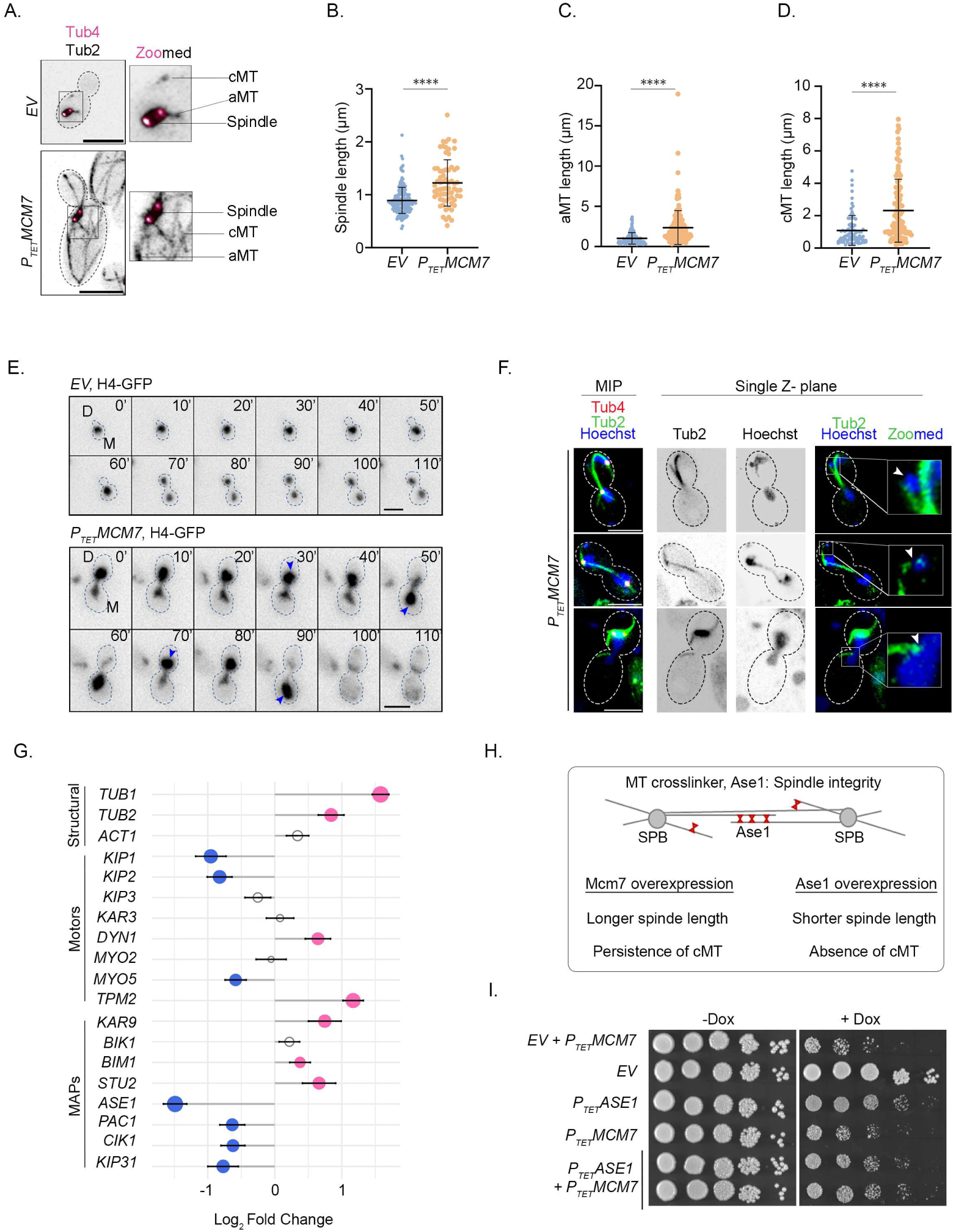
Changes in the expression of microtubule regulators impact the microtubule landscape, resulting in aberrant nuclear movement upon Mcm7 overexpression. A) Visualization of MTs by Tub2-GFP (β-tubulin) and SPB by Tub4-mCherry in *EV* (CaRG080) and *P_TET_MCM7* (SRICa157). Quantification of MT-length for B) spindle, C) aMT and D) cMT, in *EV* and *P_TET_MCM7* treated with Dox for 8 h. *n* ≥ 100 cells, unpaired *t*-test, *P*-value: (****) < 0.0001. Error bars indicate <u>s</u>tandard <u>d</u>eviation (SD). E) Live-cell microscopy montages of H4-GFP to chase nuclear dynamics in (*top*) *EV* (SRICa175) and (*bottom*) *P_TET_MCM7* (SRICa178) treated with Dox. Blue arrowheads indicate unusual nuclear oscillation. The top corner represents time in min. Scale bar: 5 µm. F) Fluorescence microscopy highlighting instances of association of aMTs/cMTs with part of chromatin moving away from the bulk chromatin, indicated by white arrowhead. MT is visualized by Tub2-GFP, chromatin is stained with Hoechst, and Tub4-mCherry marks the SPB. MIP: Maximum intensity projection. A single Z-plane representation is used for Tub2-Hoechst contact to avoid false associations caused by MIP across Z-slices. Scale bar: 5 µm. G) Lollipop graph generated from RNA-seq experiment, depicting fold-change in the expression of major microtubule-associated components in *P_TET_MCM* cells compared to *EV* control. Lollipops at the positive *x*-axis or with pink heads are up-regulated, whereas lollipops with blue heads or towards the negative *x*-axis are down-regulated. Non-colored lollipop heads are non-significant. H) (*Top)* Schematic representation of Ase1 function, transcripts of which are downregulated upon Mcm7 overexpression. (*Below)* Table comparing contrasting MT phenotypes upon overexpression of Ase1 and Mcm7 individually. I) Ten-fold serial dilution spotting assay to assess growth differences in Mcm7 overexpression (*P_TET_MCM7*-CaPJ165) and Mcm7-Ase1 co-overexpression (*P_TET_ASE1*+ *P_TET_MCM7*-SRICa225). Strain harboring both *P_TET_MCM7* and *EV* (SRICa220) serves as a negative control. Plates were imaged after 48 h of incubation at 30 °C.

The MT network is carefully orchestrated to ensure proper nuclear positioning and movement during chromosome segregation (Varshney and Sanyal 2019). Given the extensive alterations in MT architecture observed upon Mcm7 overexpression, we next examined whether these defects impacted chromatin behavior and nuclear dynamics. Whereas control cells segregated their nuclei at the bud neck, Mcm7-overexpressing cells often displayed nuclear oscillations, characterized by repeated back-and-forth movement of the nucleus between mother and daughter cells (Fig. 3E, blue arrowheads). We frequently observed aMTs and cMTs associated with chromatin fragments or chromatin regions detached from the main nuclear mass, which might explain the defective chromatin movement and positioning (Fig. 3F). These results indicate that elevated Mcm7 levels disrupt MT-dependent nuclear positioning and movement, providing a mechanistic basis for the chromosome segregation defects and genome instability associated with Mcm7 overexpression.

### Overexpressed Mcm7-induced transcriptional reprogramming perturbs microtubule homeostasis

MT properties are regulated by the structural components (α- and β-tubulin), the motor proteins of the kinesin and dynein families and the <u>m</u>icrotubule-<u>a</u>ssociated <u>p</u>roteins (MAPs) (Hildebrandt and Hoyt 2000; Meier et al. 2024). While there are no reports supporting a direct role of Mcm7 in MT dynamics, its dysregulation may influence the MT network through alterations in chromatin organization or transcriptional programming, given our observation of major transcriptional alterations. RNA-seq analysis revealed elevated levels of both α- and β-tubulin transcripts, while actin levels remained unchanged (Fig. 3G). Motor protein expression was also perturbed, with reduced *KIP1* and *KIP2* and increased dynein (*DYN1*) transcripts (Fig. 3G). Among MAPs, *BIM1*, *KAR9*, and *STU2* were modestly upregulated (Fig. 3G). Notably, *ASE1*, encoding a spindle midzone protein that stabilizes antiparallel interpolar MTs through crosslinking, was downregulated (Fig. 3G-H). Loss of Ase1 function results in bent, fragile spindles during elongation in the fission yeast *Schizosaccharomyces pombe* (Yamashita et al. 2005), resembling the spindle abnormalities observed in our study. Overexpression of *ASE1* in *C. albicans* (SRICa236) produced phenotypes distinct from those of Mcm7 overexpression, including loss of cMTs at stages when they are normally present and a higher proportion of cells with shorter spindles (Fig. S4D–E), in contrast to the elongated spindles and persistent cMTs seen upon Mcm7 overexpression (Fig. 3H). The opposing consequences of increasing Ase1 or Mcm7 dosage provided the rationale for a dosage-compensation approach, in which we tested whether co-overexpression of *ASE1* could ameliorate the cell lethality induced by excess Mcm7. Indeed, the strain co-overexpressing Ase1-Mcm7 (SRICa225) exhibited a modest improvement in growth compared to Mcm7 overexpression alone (Fig. 3I), suggesting that partial modulation of MT-related phenotypes may mitigate certain aspects of the Mcm7-induced defects. However, the incomplete rescue of growth suggests that the defects caused by Mcm7 overexpression likely arise from perturbations in a broader protein network beyond Ase1. In addition, the intrinsic growth defect associated with Ase1 overexpression may limit the extent of phenotypic restoration (Fig. 3I). Together, these findings establish a functional genetic interaction between Mcm7 dosage and the MT regulatory network, highlighting how an imbalance in replication factor levels can influence MT architecture and mechanics.

### Excess Mcm7 accumulates in the nucleolus and perturbs nucleolar homeostasis

Our results, described above, indicate that Mcm7 levels play a role in maintaining proper expression profiles of MT modulators. However, Mcm7 has been shown to physically interact with the motor protein dynein in bladder cancer stem-like cells (Mo et al. 2022). Immunofluorescence assays in the human hepatoblastoma HepG2 cell line revealed colocalization of Mcm7 with β-tubulin after nuclear envelope breakdown (Zheng et al. 2017). While MCMs are known to be nuclear proteins, information regarding their distribution upon overexpression remains scarce. Hence, we investigated the localization pattern of Mcm7 under both native and overexpression conditions in *C. albicans*. We functionally expressed C-terminally tagged Mcm7 with fluorescent reporters (mCherry-SRICa075 and SRICa082/GFP-SRICa239 and SRICa244) and confirmed its expression by western blotting (Fig. 4A). Mcm7-GFP under native conditions (SRICa244) showed a stage-specific co-localization with Hoechst-stained chromatin, as described for most MCM subunits across species (Young and Tye 1997a; Sreekumar et al. 2021) (Fig. 4B), with loss of signals in the large-budded cells with an unsegregated nuclear mass. Unexpectedly, upon overexpression (SRICa239), Mcm7 showed two types of localization anomalies: a) in contrast to wild-type, we observed persistence of Mcm7 in the nucleus at G2/M stage with an unsegregated nucleus, and b) strong localization signals at the proximity of the Hoechst-stained chromatin, resembling the nucleolus (Fig. 4C-D). Indeed, co-localization of Mcm7-GFP with Nop1-mCherry (SRICa243), a nucleolar protein (Reza et al. 2024), confirmed that Mcm7 is ectopically localized to the nucleolus upon overexpression (Fig. 4D). Importantly, this altered distribution was detectable as early as 4 h after induction and was present at all stages of the cell cycle, indicating that nucleolar localization of Mcm7 is specific and not merely a secondary consequence of prolonged cell cycle arrest or aberrant morphology (Fig. 4D).

**Figure 4.**
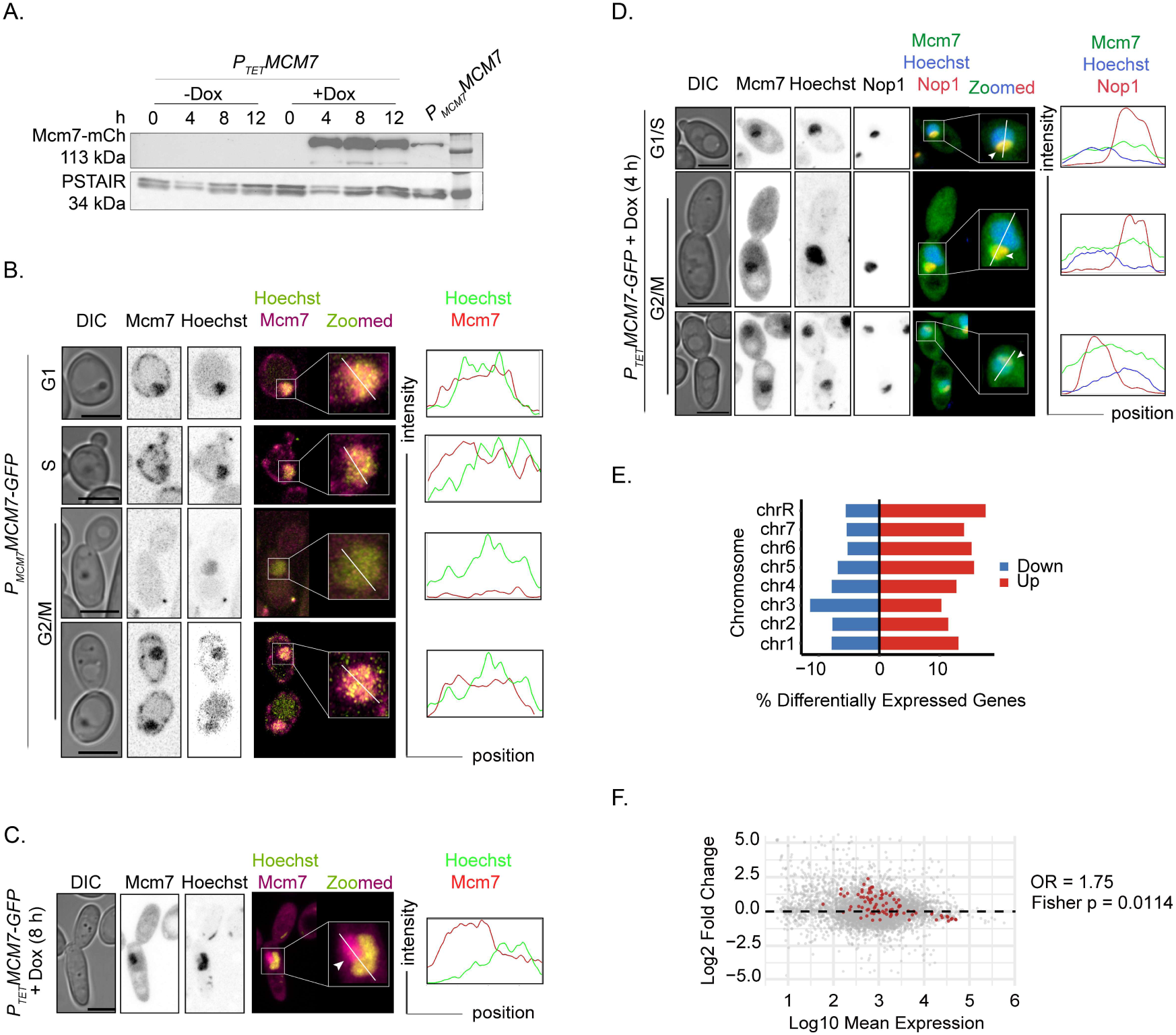
Excess Mcm7 shows ectopic nucleolar accumulation and upregulates the transcription of genes associated with nucleolar function. A) Western blotting confirmation of fluorescence epitope-tagged Mcm7-mCherry expressed from native promoter (*P_MCM7_MCM7-* SRICa075) and from Dox-inducible tetracycline promoter for overexpression (*P_TET_MCM7*-SRICa082). B) Fluorescence microscopy images showing cell cycle-dependent nuclear localization of Mcm7-GFP expressed from its native promoter (SRICa244). Nucleus is stained with Hoechst. Scale bar: 5 µm. C) Fluorescence microscopy images of Mcm7-GFP expressed from the *P_TET_* overexpression promoter (SRICa239) in the presence of Dox for 8 h. The white arrowhead indicates localization outside the Hoechst-stained nucleus. Scale bar: 5 µm. D) Fluorescence microscopy and line scan plot profile to map the co-localization pattern of Mcm7-GFP with nucleolus marked by Nop1-mCherry (SRICa243) and chromatin stained with Hoechst in different stages of the cell cycle after induction with Dox for 4 h. The white arrowheads indicate nucleolar localization of Mcm7. Scale bar: 5 µm. E) Bar plot representing the percentage of <u>d</u>ifferentially <u>e</u>xpressed <u>g</u>enes (DEGs) from each chromosome, normalized to their respective total number of genes. F) A <u>m</u>inus-<u>a</u>verage (MA) plot of DEGs (grey spots) overlaid with genes associated with nucleolus and ribosome biogenesis (a total of 92 genes obtained from Candida Genome Database) as maroon spots. Statistical significance: Fisher’s test *p*-value 0.0114. Odds ratio (OR) value interpretation: >1 means gene set is upregulated or over-represented, =1 means no association, <1 means gene set is downregulated or under-represented amongst the DEGs.

The nucleolus is a phase-separated nuclear compartment, critical for ribosome biogenesis, RNA metabolism, and cellular stress response (Dubois and Boisvert 2016). Since elevated Mcm7 levels are associated with transcriptional changes, we examined the consequence of such unexpected accumulation of the protein on nucleolar-associated genes. We observed that the highest percentage of upregulated genes, normalized to gene number per chromosome, was in chromosome R, which harbors the rDNA loci residing in the nucleolus (Fig. 4E). Overlaying nucleolar and ribosome biogenesis genes onto the RNA-seq dataset revealed a significant skew toward upregulation (Fisher’s exact test, *p* = 0.0114) (Fig. 4F). Although alterations in ribosomal gene expression can arise during general cellular stress responses, the spatial relocalization of Mcm7 to the nucleolus raises the possibility that excess Mcm7 directly perturbs nucleolar function.

This study in *C. albicans* identifies a critical role for Mcm7 dosage in regulating chromosome segregation and nucleolar function, likely through transcriptional rewiring. To extend these findings across cancer types, we analyzed gene co-expression patterns using cBioPortal for Cancer Genomics (https://www.cbioportal.org/), with data from The Cancer Genome Atlas and International Cancer Genome Consortium. To briefly summarize the analysis pipeline (Fig. 5A), genes with a Spearman’s correlation coefficient ≥ 0.4 (implicating positive correlation in expression) for each MCM subunit were selected, and a set intersection analysis was performed, wherein we obtained lists of genes unique to each MCM subunit as well as the overlapping genes in all possible combinations (Fig. S5A). The unique gene sets were then subjected to <u>g</u>ene <u>o</u>ntology (GO) analysis. For MCM4, MCM5 and MCM6, biological processes pertaining to development, immune response and cellular response to external stimuli were enriched, respectively (Fig. S5B). GO enrichment analysis for MCM2 and MCM3 did not produce enrichment plots, likely due to the small number of genes in their respective correlated gene sets (Fig. S5A). Notably, genes uniquely associated with MCM7 were significantly enriched for ribosome biogenesis and RNA processing pathways (Fig. 5B), closely mirroring the transcriptional signature observed following Mcm7 overexpression in *C. albicans*.

**Figure 5.**
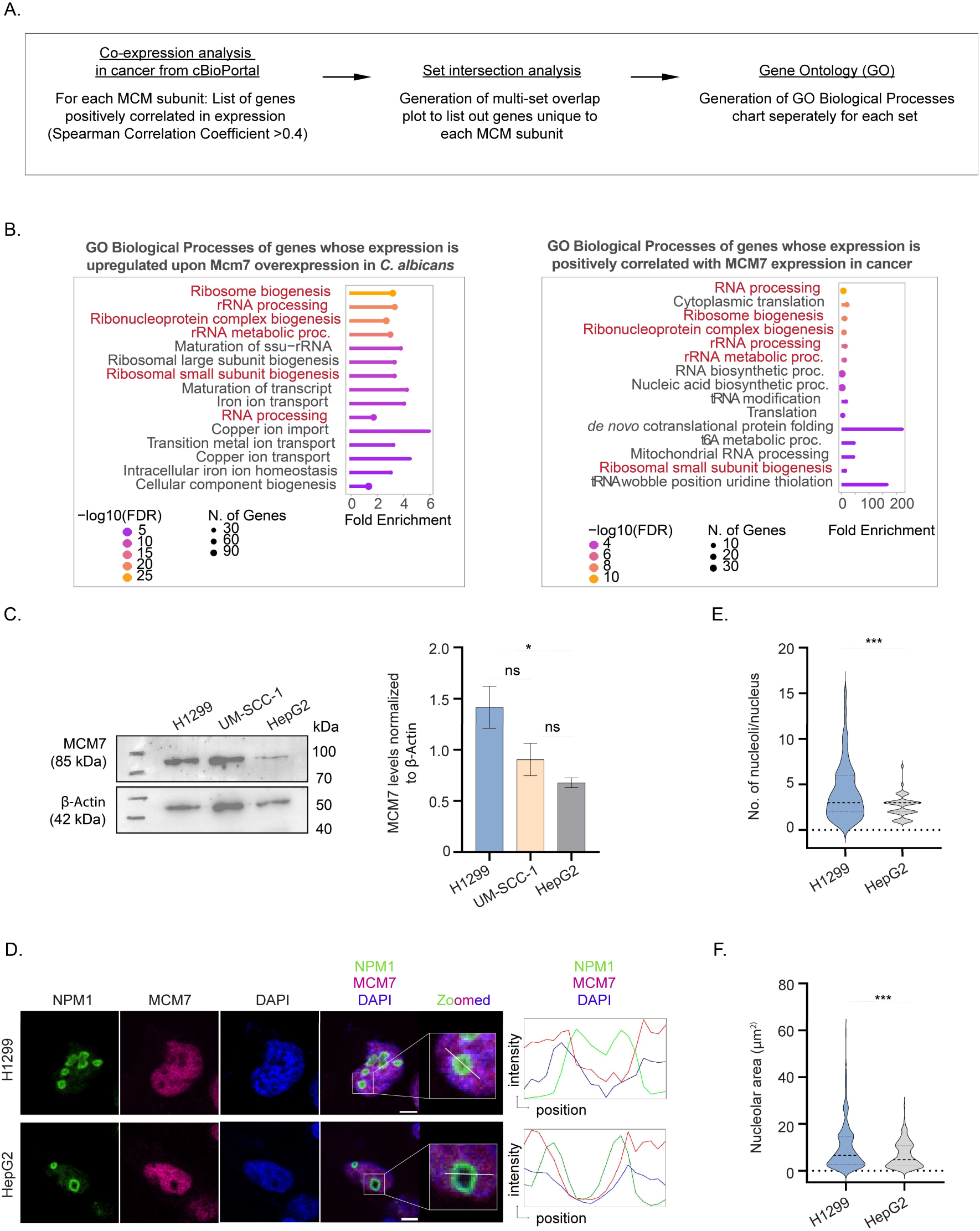
Mcm7 dosage is closely associated with nucleolar programs from yeast to humans. A) Overview of the bioinformatic workflow used to identify biological processes associated with dysregulated MCM subunit expression in cancer. B) Comparison of <u>G</u>ene <u>O</u>ntology terms for <u>B</u>iological <u>P</u>rocesses (GO-BP), for (*left*) the list of genes upregulated upon Mcm7 overexpression in *C. albicans* obtained from RNA-seq experiment, and (*right*) the list of genes whose expression is uniquely positively correlated to MCM7 expression across several cancer types, obtained from cBioPortal. The common GO-BP terms across both sets are highlighted in maroon. GO analysis was conducted in ShinyGO 0.85.1 for both species. C) (*left*) Western blot and (*right*) quantification to measure cellular levels of MCM7 in three different human cell lines. β-Actin levels were used for normalization. *N* =3, Ordinary One-way ANOVA with uncorrected Fisher’s LSD. (*) P-value <0.05, (ns) > 0.05. D) Indirect immunofluorescence assay for sub-cellular localization of MCM7 with respect to the nucleoli marked by NPM1 and nuclear chromatin stained by DAPI. Scale bar: 5 µm. E) Quantification of the number of nucleoli per nucleus across cell lines. *n* ≥ 50 cells, unpaired t-test. (***) P-value <0.001. F) Quantification of the area of individual nucleoli across cell lines. *n* ≥ 150 nucleoli, unpaired t-test. (***) P-value <0.001.

To investigate whether these observations are also reflected in human cells, we next examined endogenous MCM7 expression in three human cancer cell lines: the non-small cell lung carcinoma cell line (H1299), the head and neck squamous cell carcinoma cell line (UM-SCC-1), and the hepatocellular carcinoma cell line (HepG2). Immunoblot analysis revealed a gradient of MCM7 expression across these cell lines, with the highest levels observed in H1299 and the lowest levels in HepG2 (Fig. 5C). The marked difference in MCM7 expression between H1299 and HepG2 provides an opportunity to assess whether variation in MCM7 abundance is associated with differences in nucleolar organization. Since nucleolar size and number are widely used as morphological indicators of nucleolar activity and ribosome biogenesis, and are frequently altered in cancer (Montanaro et al. 2008; Dubois and Boisvert 2016), we next compared these parameters between the two cell lines. Immunofluorescence microscopy showed that MCM7 localized predominantly to the interphase nucleus and largely overlapped with DAPI-stained chromatin in both H1299 and HepG2 cells, with no detectable nucleolar enrichment (a phenomenon observed upon Mcm7 overexpression in *C. albicans*) (Fig. 5D). Intriguingly, H1299 cells contained more nucleoli per nucleus than HepG2 cells (Fig. 5E). Furthermore, nucleoli were significantly larger in H1299 cells (Fig. 5F). The enrichment of ribosome biogenesis and RNA processing pathways among MCM7-associated genes in pan-cancer transcriptomic datasets, coupled with the larger and more abundant nucleoli observed in cells expressing higher levels of MCM7, hints at a broader role for MCM7 dosage in coordinating nucleolar homeostasis across evolutionarily distant systems.

## Discussion

In this study, we identify Mcm7 as a dosage-sensitive regulator of genome stability whose overexpression, unlike that of other pre-RC subunits, is sufficient to induce CIN and cell death. We demonstrate that excess Mcm7 disrupts its normal cell cycle-dependent localization, preventing its efficient nuclear exclusion during G2/M and promoting its unexpected accumulation within the nucleolus, a subnuclear localization not previously described for any MCM subunit (Fig. 6A-B). This perturbation is accompanied by widespread transcriptional reprogramming that affects not only the transcription of genes associated with nucleolar function but also the expression of genes involved in MT homeostasis, leading to downstream consequences such as disruption of spindle integrity, altered MT length, defective centromere organization and aberrant nuclear movement. This cascade of defects ultimately leads to impaired chromosome segregation and SAC-mediated cell cycle arrest, culminating in cell death (Fig. 6A). Moreover, the convergence of pan-cancer transcriptomic analyses and human cell-based observations, both linking MCM7 expression to nucleolar biology, extends the relevance of our findings beyond *C. albicans* (Fig. 6B). Together, this study reveals an unanticipated role for Mcm7 dosage in coordinating nucleolar and cytoskeletal homeostasis to preserve genome stability, thereby expanding the functional repertoire of this conserved replication factor.

**Figure 6.**
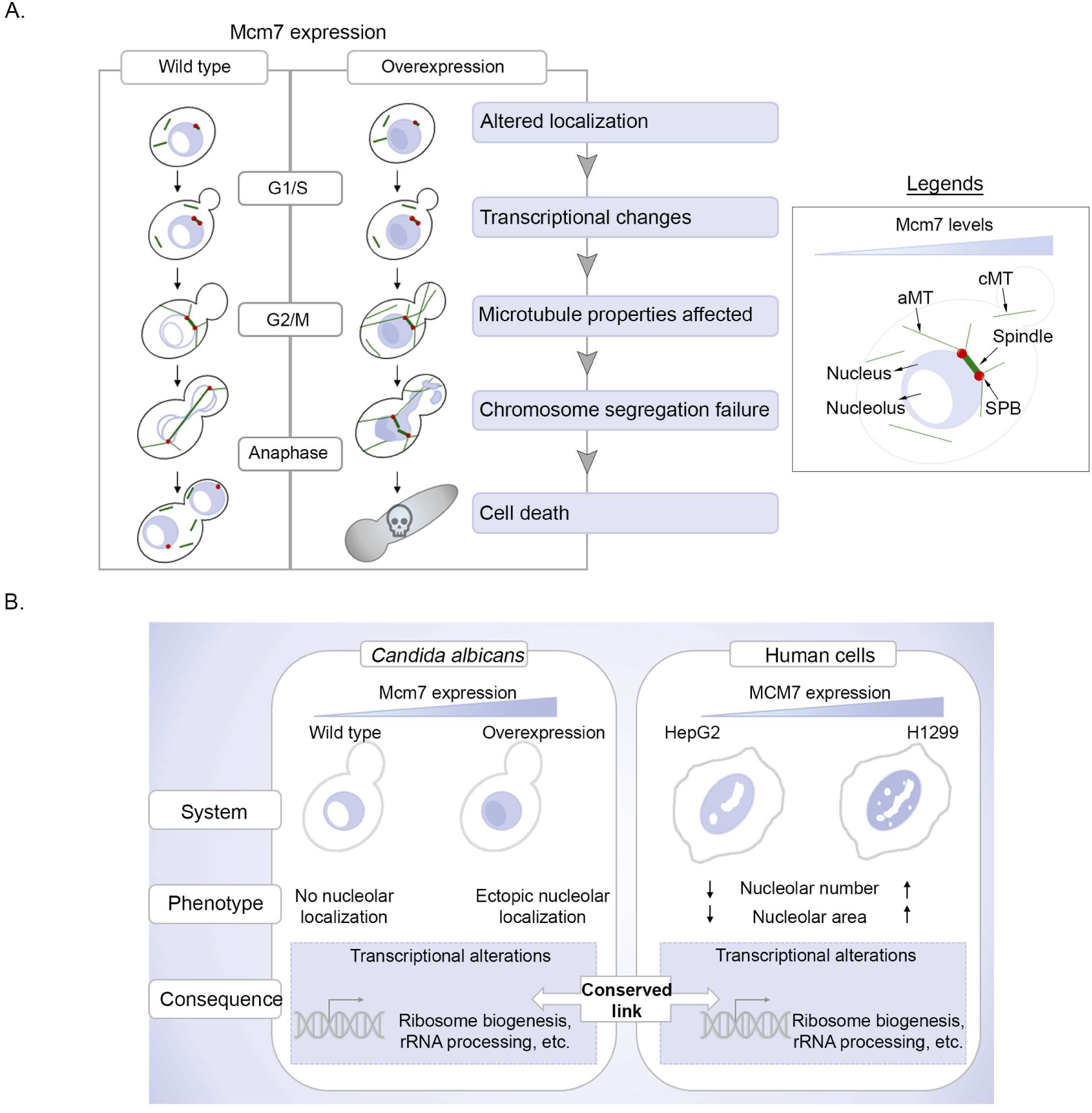
Mcm7 dosage coordinates microtubule and nucleolar homeostasis. A) Schematic encapsulating our proposed framework for Mcm7 dosage-dependent regulation of cellular homeostasis. Under wild-type conditions in *C. albicans*, nuclear localization of Mcm7 is cell cycle stage-specific. Elevated Mcm7 levels lead to aberrant nucleolar accumulation and altered expression of genes involved in ribosome biogenesis and RNA processing. Concurrently, Mcm7 is retained in the nucleus even at the G2/M stage, contributing to widespread transcriptional changes, including the genes involved in the MT machinery. MT homeostasis is disrupted, resulting in increased aMT and cMT length and spindle defects, which in turn lead to aberrant centromere clustering and nuclear dynamics. These defects compromise chromatin stability, activate the SAC, and culminate in cell cycle arrest and cell death. B) Comparative summary of the relationship between Mcm7 dosage and nucleolar homeostasis in *C. albicans* and human cancer cells. In *C. albicans*, Mcm7 overexpression promotes ectopic nucleolar localization accompanied by transcriptional upregulation of ribosome biogenesis, rRNA processing, and related nucleolar pathways. In human cancer cell lines, increasing endogenous MCM7 expression (from HepG2 to H1299) correlates with an increase in the number and size of nucleoli. Furthermore, enrichment of the same nucleolar-associated transcriptional programs is observed in cancer cells and tissues with dysregulated MCM7 levels, suggesting a conserved association.

MCMs are cyclic proteins, both in terms of their expression and nuclear localization/chromatin binding. Nuclear exclusion, particularly during the mitotic phase of the cell cycle, is observed for multiple MCM subunits across species, including *S. cerevisiae* (Dalton and Whitbread 1995; Young and Tye 1997b), plants (Shultz et al. 2009) and human cell lines (Méndez and Stillman 2000). A comparable localization pattern follows in *C. albicans*, wherein nuclear exclusion is observed at the G2/M stage prior to nuclear division for both Mcm2 (Sreekumar et al. 2021) and Mcm7. Thus, a stringent regulation of cell cycle dependent chromatin association for MCMs is evolutionarily established, likely as a mechanism to prevent re-replication. It may additionally be essential for restricting other deleterious consequences associated with the diverse cellular functions of MCM proteins. A striking outcome of Mcm7 overexpression in *C. albicans* is its aberrant retention in the nucleus during G2/M together with its unexpected accumulation in the nucleolus. This wrong-place-and-the-wrong-time scenario has significant implications for the cell’s transcriptional response, as evidenced by marked changes in the expression of genes implicated in nucleolar function. The nucleolus is a phase-separated module, and components of the pre-RC in organisms such as *S. cerevisiae*, *Drosophila melanogaster* and humans exhibit liquid-liquid phase separation (Parker et al. 2019; Hossain et al. 2021; Wu et al. 2024). It is possible that overexpression of Mcm7 in *C. albicans* perturbs its nuclear concentration, driving phase separation of the protein with the nucleolus and leading to its ectopic localization. Alternatively, overexpression may alter its interaction network, forming new associations with nucleolar proteins. Delineating the interacting partners of Mcm7 under native and over-expressed conditions will provide further insights.

MCM7 has been shown to act as a transcriptional co-factor for the <u>a</u>ndrogen receptor (AR) in human prostate cancer cell lines. This association is critical in regulating the expression of AR-responsive genes. Notably, this association also facilitates the self-regulation of MCM7 (Shi et al. 2008). Such self-regulation of expression was also reported in *S. cerevisiae*, wherein Mcm7 acts as a transcriptional co-factor for Mcm1, modulating its own expression and the transcription of early cell cycle genes, including many pre-RC proteins (Fitch et al. 2003). It seems there exists a strong auto-regulatory circuit for Mcm7, which is likely to be perturbed upon overexpression. In *C. albicans,* Mcm1 regulates the expression of cyclic genes involved in spindle function, including Kip2 and Ase1 (Côte et al. 2009), both of which are downregulated upon Mcm7 overexpression. Our dosage-compensation genetic assay with Ase1 and Mcm7 suggests that the spectrum of MT phenotypes arises from dysregulated expression of cytoskeletal components sensitive to Mcm7 levels. The ectopic localization of Mcm7 to the nucleolus, coupled with increased transcription of nucleolus-associated genes, further supports the idea that Mcm7 may act as a transcriptional modulator, affecting a critical network of proteins controlling cellular homeostasis.

Although this study establishes a novel association among multiple seemingly unrelated pathways, a key limitation is that it does not identify a direct molecular pathway linking nucleolar perturbation to MT defects. While genetic, cytological, and transcriptomic analyses reveal a strong association between these processes via Mcm7, resolving the intermediates that connect them is particularly important, as the observations may extend beyond fungal biology. Morphological and functional changes in the nucleolus are well-established hallmarks of cancer, owing to its indispensable role in ribosome biogenesis, gene expression and protein synthesis, cellular senescence, and the stress response (Montanaro et al. 2008; Orsolic et al. 2016; Hwang and Denicourt 2024). The unique association of MCM7 expression with genes involved in nucleolar function across cancer types, along with a possible correlation between nucleolar morphology and MCM7 expression in human cell lines, provides a strong rationale for future investigation (Fig. 6B). Similarly, the roles of motors and MAPs in cancer metastasis are emerging due to their importance in mediating DNA repair, reorganizing MTs during morphological transitions associated with cell movement and in the translocation of metastatic signaling molecules (Parker et al. 2014; Wattanathamsan and Pongrakhananon 2022; Mangaonkar et al. 2024). In this context, our study establishes a functional link between dysregulated Mcm7 expression and concomitant perturbations in both nucleolar biology and MT properties, an association that can be examined and exploited to better understand oncogenesis and to design combinatorial cancer therapies.

## Materials and Methods

### Strains, plasmids, and primers

Strains, plasmids, primers and human cell lines used in this study are provided in Supplementary tables S1, S2, S3 and S4, respectively. Chemicals and antibodies used in the study are listed in Supplementary Table S5. Cloning and strain construction strategies are outlined in the supplementary methods.

### Media and growth conditions

*C. albicans* cells were grown at 30 °C in YPDU (1% yeast extract, 2% peptone, 2% dextrose, 0.1 µg/mL uridine) or complete medium (CM, 2% dextrose, 1% yeast nitrogen base and auxotrophic supplements with amino acids histidine, arginine, leucine, uridine, adenine sulfate, and tryptophan (0.1 µg/mL)). Agar (2%) was added to the solid medium preparation. Nourseothricin (NAT) and hygromycin B (Hyg B) were used at final concentrations of 100 µg/mL and 800 µg/mL, respectively, in YPDU medium. Anhydrotetracycline (Atc, 3 µg/mL) or doxycycline (Dox, 50 µg/mL) was added to YPDU media to induce overexpression of genes from the tetracycline-inducible promoter (*P_TET_*). Cultures were grown at 30 °C in the dark, as these compounds are light sensitive. For depletion of protein from the conditional repression promoter (*P_MET3_*), cells were grown at 30 °C either in permissive (YPDU/CM) or non-permissive (YPDU/CM + 5 mM methionine (M) + 5 mM cysteine (C)) conditions.

*E. coli* strains containing the CIp10-*P_TET_*-GTW derivatives were grown at 30 °C with ampicillin (100 µg/mL) selection. All other *E. coli* strains were grown at 37 °C with ampicillin (100 µg/mL) or chloramphenicol (34 µg/ml), depending on the selection marker. Chemically competent *E. coli* cells were prepared as described in (Chung et al. 1989).

### Human Cell Culture

Human lung carcinoma H1299 cell line (ATCC) was cultured in RPMI 1640 media supplemented with 2 mM glutamine and 10% (v/v) fetal bovine serum (FBS). Human Hepatocellular Carcinoma HepG2 cell line (ATCC) was grown in Dulbecco’s modified Eagle’s medium (DMEM), supplemented with 10% (v/v) FBS. The human Head and Neck Squamous cell carcinoma UM-SCC-1 cell line was grown in Dulbecco’s Modified Eagle Medium/Nutrient Mixture F-12 Ham (DMEM/F12, 1:1 mixture) supplemented with 10% (v/v) FBS. Proliferating cell cultures were maintained in a 5% CO_2_ humidified incubator at 37 °C and 90% relative humidity.

### Dilution spot assay

Cells equivalent to OD_600_ of 1/mL were pelleted from an overnight culture in YPDU, washed and resuspended in 1 mL water, serially diluted 1:10, and 3 µL was spotted on YPDU or YPDU + Dox (50 µg/mL) agar plates as indicated.

### Flow cytometry analysis

Cells were grown overnight and transferred to fresh media with Dox the next day, with a starting OD_600_ of 0.2/mL. Cells were collected before incubation at 30 °C (0 h) and again after 8 h. Cells were fixed with 70% ethanol and treated with RNase overnight. Cell suspension was stained with Propidium iodide, briefly sonicated and subjected to flow cytometry (FACSAria III, BD Biosciences). A minimum of 30000 events were captured, and the outputs were analyzed using the FlowJo software.

### Cell lysis and western blotting

For *C. albicans*, cells were collected at an OD_600_ of 3.0 and precipitated with 16% Trichloroacetic acid (TCA) overnight at -20 °C. The pellets were then washed twice with chilled 80% acetone and resuspended in lysis buffer (1% SDS, 1N NaOH) with SDS loading dye. Samples were boiled for 10 min and electrophoresed on a 10% or 12% polyacrylamide gel. Following protein transfer on a nitrocellulose membrane using either the semi-dry or wet-transfer method (wet-transfer method used for Mcm7-mCherry western), the blot was blocked with 5% skimmed milk in 1x PBS for an hour. Post-blocking, primary antibody incubation was performed overnight at 4 °C (rabbit anti-Protein A, mouse anti-PSTAIR, rat anti-RFP, and rabbit anti-H3, all used at 1:5000 dilution in 2.5% skimmed milk in 1x PBS). The following day, blots were washed 3 times in 1x PBS-Tween (0.05%) and incubated with secondary antibody for 1 h (goat anti-rabbit IgG-HRP, goat anti-mouse IgG-HRP, rabbit anti-rat IgG-HRP, used at a dilution of 1:10,000 in 2.5% skimmed milk). Following three PBST washes, the blot was developed using the chemiluminescence method. Quantification was done using FIJI ImageJ.

For mammalian cell lines, cells were washed twice with PBS and lysed on ice in RIPA buffer (50 mM Tris-HCl, pH 7.4, 150 mM NaCl, 1% NP-40, 0.5% sodium deoxycholate, and 0.1% SDS) supplemented with protease inhibitor cocktail. Lysates were incubated on ice for 10 min with intermittent mixing and clarified by centrifugation at 12,000× g for 20 min at 4°C. The supernatants were collected, and protein concentrations were determined using the Bradford assay. Equal amounts of protein (15 μg) were resolved on SDS-polyacrylamide gels and transferred onto PVDF membranes. Membranes were blocked with 3% w/v non-fat skim milk in TBS containing 0.1% Tween-20 (TBST) for 1 h at room temperature and thereafter incubated overnight at 4°C with primary antibodies (rabbit anti-β-Actin (1:10,000), rabbit anti-MCM7 (1:1,000) prepared in 3% Skim milk in 1X TBST). The following day, the blots were washed three times with 1X PBS, and membranes were incubated with HRP-conjugated secondary antibodies (goat anti-rabbit IgG-HRP (1:10,000)) for 1 h at room temperature. Immunoreactive bands were visualised using enhanced chemiluminescence (ECL) reagents and imaged using a chemiluminescence detection system. β-Actin was used as the loading control. Quantification was done using FIJI ImageJ.

### Chromatin immunoprecipitation

For CENPA^Cse4^-TAP ChIP, cells were collected at an OD_600_ of 100 and cross-linked with formaldehyde to a final concentration of 1% for 15 min. The ChIP protocol was followed as described previously (Sreekumar et al. 2021). Briefly, cells were spheroplasted with Zymolyase-20T, and the chromatin was sheared using Bioruptor Pico from Diagenode. The sheared chromatin was split into three fractions: Input (-ab, -beads), +Ab (+Ab, +beads), and -Ab (-Ab, +beads). Incubation with anti-Protein A antibody was performed overnight at 4 °C with constant rotation. The following day, Protein A-Sepharose beads were added to +Ab and -Ab fractions and incubated at 4 °C with constant rotation. Post-washing, the samples were de-crosslinked, purified, and ethanol-precipitated. The pellets were resuspended in Milli-Q water and subjected to downstream PCR reactions. All ChIP-qPCR experiments were performed with three technical replicates for each sample. Enrichment was calculated by the % input method (Sreekumar et al. 2021).

### Sample preparation for fluorescence microscopy, image acquisition, and quantification

For all microscopy experiments involving Dox treatment, cultures were grown overnight in YPDU, and cells were transferred to fresh YPDU media the next day with or without treatment (starting OD_600_ of 0.2/mL). Cells were collected after 8 h or at the indicated time points and washed once/twice with water. Hoechst dye was added to the cell suspension before imaging to visualize chromatin, wherever mentioned. For replication poison experiments, overnight cultures were secondary inoculated into fresh YPDU medium the next day, starting at an OD_600_ of 0.2/mL and grown until OD_600_ reached 1/mL. 0.005% methylmethane sulfonate (MMS) or 10 mM hydroxyurea (HU) was then added to the secondary cultures, which were then grown for the indicated time points. Cells were collected for FACS, and the remaining cells were pelleted, washed, and imaged after Hoechst addition. For live-cell imaging, cells were adhered to Concanavalin A-treated glass-bottomed round dishes and supplemented with CM + Dox media. A temperature of 30 °C was maintained during image acquisition.

For indirect immunofluorescence in mammalian cells, H1299 and HepG2 cells were grown on poly-L-lysine-coated cover slips at 37 °C in a 5% CO2 incubator. The media was removed, and the cell layer was washed in PBS to ensure all the media was removed. The cells were fixed in 4% paraformaldehyde (PFA) for 10 min at room temperature. The cells were washed with PBS to remove the remnant PFA. The cells were then permeabilized in 1% Triton X-100. Cells were washed in PBS to remove the residual TritonX-100. The cells were blocked in blocking solution (5% FBS in PBS) at 37 °C for 45 min. The blocked cells were then incubated in primary antibody (mouse anti-NPM1 (1:500) and rabbit anti-MCM7 (1:200)) for 1 h at room temperature on a reciprocal shaker. The cells were washed in washing buffer (1% FBS in PBS). The cells were then incubated in Alexa-fluor conjugated secondary antibody (Alexa 488 and Alexa 568) corresponding to the primary antibody used, and incubated at room temperature for 1 h. The nucleus was counter-stained with 0.1 μg/ml of DAPI in PBS for 5 min to visualize the nuclei. Excess DAPI stain was washed off with PBS. The cover slips were mounted in 70% glycerol onto a glass slide.

Imaging was performed either on an LSM880 Airyfast confocal microscope using a Plan-Apochromat 63×/1.4 Oil DIC M27/100×/1.4 Oil DIC M27 objectives or using a Zeiss Axio Observer 7, 100×/1.4 Plan Apochromat, pco. edge 4.2 sCMOS. All images were processed using FIJI ImageJ software.

Spindle length, astral and cytoplasmic microtubules were quantified using a straight line or freehand (for curved or bent microtubules) tool in FIJI ImageJ software after maximum intensity projection of the image. Nucleolar area was quantified by freehand outline of each nucleolus after average intensity projection of the image in FIJI ImageJ software. Plot profiles were generated using the plot profile feature in FIJI ImageJ on a single Z-plane for each channel.

### RNA isolation and quantification by real-time PCR

Cells equivalent to OD_600_ of 10 were resuspended in 400 µL of lysis buffer (2.5 M sorbitol, 0.5 M EDTA, Zymolyase-20T, and β-mercaptoethanol) and then spherolasted using Zymolyase-20T at 30 °C for 1 h. The total RNA was isolated using TRIzol reagent. 500 µL of TRIzol was added to the spheroplasted cells, mixed, and incubated for 5 min at room temperature. To this, 200 µL of chloroform was added, and the solution was vortexed for 30 sec. After a 5 min incubation at room temperature, the cells were centrifuged for 15 min at 13,000 rpm at 4 °C. The aqueous phase was collected in a fresh tube and treated once again with TRIzol and chloroform. After centrifugation, 500 µL of isopropanol was added to the aqueous phase, which was incubated at room temperature for 10 min, and the RNA pellet was precipitated. The total RNA pellet was washed twice with 70% ethanol, and the air-dried pellet was resuspended in autoclaved milliQ water. 2 µg of DNase-treated RNA were taken forward for 1^st^ strand synthesis using the manufacturer’s protocol (RevertAid First Strand cDNA Synthesis Kit, ThermoFisher Scientific). Quantitative PCR (qPCR) was performed using SYBR green dye. All qPCRs were performed with three technical replicates for each sample. The relative abundance of target gene transcripts was normalized to that of actin (*ACT1*), and the fold change was calculated using the 2^-ΔΔCt^ method.

### Transcriptome analysis

RNA-seq library preparation and sequencing were outsourced from the National Centre for Biological Sciences (NCBS). Details include a) mRNA isolation was done using NEBNext Poly(A) mRNA Magnetic Isolation Module, and b) RNA library preparation was done using NEBNext® Ultra™ II Directional RNA Library Prep with Sample Purification Beads. Samples were sequenced on NovaSeq 6000 platform. Raw paired-end reads were quality-checked and adapter-trimmed using Trim Galore with FastQC. Trimmed reads were aligned to the reference genome using STAR. Alignments were generated as coordinate-sorted BAM files and indexed with SAMtools. Gene-level read counts were obtained with HTSeq-count using the corresponding GFF annotation and gene IDs as features. Differential gene expression analysis was performed in R Studio using the DESeq2 package. All downstream analyses and visualizations were performed in R Studio using available R packages.

### Cancer data analysis

In the cBioPortal (https://www.cbioportal.org/), the pan-cancer analysis of whole genomes (ICGC/TCGA) option was selected for PanCancer studies queried by gene name. mRNA expression with a z-score threshold of ±2.0 was selected. We then downloaded a list from the co-expression analysis that included gene names and the corresponding Spearman’s correlation coefficients for each MCM gene. We manually segregated the genes with a Spearman’s correlation coefficient ≥ 0.4 (implicating positive correlation in expression), thereby generating 6 lists for each MCM subunit. Next, we provided the information to an online multiset Venn diagram generator (https://bioinformatics.psb.ugent.be/webtools/Venn/) and obtained lists of genes unique to each MCM subunit and the overlaps among them. Finally, we performed a <u>G</u>ene <u>O</u>ntology (GO) <u>B</u>iological <u>P</u>rocesses (GO-BP) analysis using ShinyGO 0.85.1 (https://bioinformatics.sdstate.edu/go/) for the gene sets unique to each MCM subunit.

### Statistical analysis

All statistical analyses were performed using GraphPad Prism 8.4.0. Details of the number of replicates and the statistical tests applied for each experiment are provided in their respective figure legends.

## Supporting information

All Supplementary data

## Data access

The RNA-sequencing data generated in this study have been submitted to the NCBI Sequence Read Archive under BioProject ID PRJNA1503116.

## Acknowledgements and Funding

We thank members of the KS Lab for their valuable suggestions throughout the study. Sincere gratitude to Rohit G. for generating the strains CaRG090 and CaRG095. We thank Dr Priya B. for generating the plasmid pFA-TAP-ARG_CIp10. We thank Tejas P. for constructing pMcm7-mCherry-ARG. We also thank Sarvleen K. for generating CIp10-*P_TET_*-MCM7-HYG. The UM-SCC-1 cell line was a kind gift from Dr Gautam Sethi (National University of Singapore) to TKK. We thank Suma B.S., Siddharth P., and Tharun L.R. at the imaging facility at JNCASR. We acknowledge Narendra N. and Aparna A. at the flow cytometry facility, JNCASR.

SD, SVB, TKK, and KS acknowledge the intramural financial support from JNCASR. Financial support from the DBT-RA Program (DBT/2020/January/58) in Biotechnology and Life Sciences is gratefully acknowledged by MHR. KS also acknowledges the financial support of the JC Bose National Fellowship (Science and Engineering Research Board, Government of India, JCB/2020/000021) and the JC Bose grant (ANRF/JBG/2025/000335/LS).

## Author contributions

Conceptualization: SD, KS; Methodology-fungal system: SD, MHR; Methodology-human cell line: SVB; Validation-fungal system: SD, MHR; Validation-human cell line: SD, SVB; Formal Analysis-fungal system: SD, MHR; Formal Analysis-human cell line: SD; Writing-original draft: SD, MHR, SVB; Writing-review and editing: SD, MHR, SVB, TKK, KS; Supervision: KS; Funding acquisition: MHR, KS.

## References

Agarwal S, Smith KP, Zhou Y, et al (2018) Cdt1 stabilizes kinetochore–microtubule attachments via an Aurora B kinase–dependent mechanism. Journal of Cell Biology 217:3446–3463. 10.1083/jcb.201705127

Bachewich C, Nantel A, Whiteway M (2005) Cell cycle arrest during S or M phase generates polarized growth via distinct signals in *Candida albicans*. Molecular Microbiology 57:942–959. 10.1111/j.1365-2958.2005.04727.x

Barton R, Gull K (1988) Variation in cytoplasmic microtubule organization and spindle length between the two forms of the dimorphic fungus *Candida albicans*. Journal of Cell Science 91:211–220. 10.1242/jcs.91.2.211

Care RS, Trevethick J, Binley KM, Sudbery PE (1999) The *MET3* promoter: a new tool for *Candida albicans* molecular genetics. Molecular Microbiology 34:792–798. 10.1046/j.1365-2958.1999.01641.x

Chauvel M, Nesseir A, Cabral V, et al (2012) A Versatile Overexpression Strategy in the Pathogenic Yeast Candida albicans: Identification of Regulators of Morphogenesis and Fitness. PLoS ONE 7:e45912. 10.1371/journal.pone.0045912

Chung CT, Niemela SL, Miller RH (1989) One-step preparation of competent Escherichia coli: transformation and storage of bacterial cells in the same solution. Proc Natl Acad Sci USA 86:2172– 2175. 10.1073/pnas.86.7.2172

Côte P, Hogues H, Whiteway M (2009) Transcriptional Analysis of the *Candida albicans* Cell Cycle. MBoC 20:3363–3373. 10.1091/mbc.e09-03-0210

Dalton S, Whitbread L (1995) Cell cycle-regulated nuclear import and export of Cdc47, a protein essential for initiation of DNA replication in budding yeast. Proc Natl Acad Sci USA 92:2514–2518. 10.1073/pnas.92.7.2514

DePamphilis ML (ed) (2006) DNA replication and human disease. Cold Spring Harbor Laboratory Press, Cold Spring Harbor, N.Y

Dubois M-L, Boisvert F-M (2016) The Nucleolus: Structure and Function. In: Bazett-Jones DP, Dellaire G (eds) The Functional Nucleus. Springer International Publishing, Cham, pp 29–49

Dutta S, Bhat K, Aggarwal R, Sanyal K (2025) Fungi as models of centromere innovation: from DNA sequence to 3-dimensional arrangement. Chromosome Res 33:18. 10.1007/s10577-025-09775-1

Finley KR, Berman J (2005) Microtubules in *Candida albicans* Hyphae Drive Nuclear Dynamics and Connect Cell Cycle Progression to Morphogenesis. Eukaryot Cell 4:1697–1711. 10.1128/EC.4.10.1697-1711.2005

Fitch MJ, Donato JJ, Tye BK (2003) Mcm7, a Subunit of the Presumptive MCM Helicase, Modulates Its Own Expression in Conjunction with Mcm1. Journal of Biological Chemistry 278:25408–25416. 10.1074/jbc.M300699200

Forsburg SL (2004) Eukaryotic MCM Proteins: Beyond Replication Initiation. Microbiol Mol Biol Rev 68:109–131. 10.1128/MMBR.68.1.109-131.2004

Gardner MK, Bouck DC, Paliulis LV, et al (2008) Chromosome Congression by Kinesin-5 Motor-Mediated Disassembly of Longer Kinetochore Microtubules. Cell 135:894–906. 10.1016/j.cell.2008.09.046

Guin K, Sreekumar L, Sanyal K (2020) Implications of the Evolutionary Trajectory of Centromeres in the Fungal Kingdom. Annu Rev Microbiol 74:835–853. 10.1146/annurev-micro-011720-122512

Hildebrandt ER, Hoyt MA (2000) Mitotic motors in Saccharomyces cerevisiae. Biochimica et Biophysica Acta (BBA) - Molecular Cell Research 1496:99–116. 10.1016/S0167-4889(00)00012-4

Honeycutt KA, Chen Z, Koster MI, et al (2006) Deregulated minichromosomal maintenance protein MCM7 contributes to oncogene driven tumorigenesis. Oncogene 25:4027–4032. 10.1038/sj.onc.1209435

Hosea R, Hillary S, Naqvi S, et al (2024) The two sides of chromosomal instability: drivers and brakes in cancer. Sig Transduct Target Ther 9:75. 10.1038/s41392-024-01767-7

Hossain M, Bhalla K, Stillman B (2021) Multiple, short protein binding motifs in ORC1 and CDC6 control the initiation of DNA replication. Molecular Cell 81:1951–1969.e6. 10.1016/j.molcel.2021.03.003

Hwang S-P, Denicourt C (2024) The impact of ribosome biogenesis in cancer: from proliferation to metastasis. NAR Cancer 6:zcae017. 10.1093/narcan/zcae017

Jaitly P, Legrand M, Das A, et al (2022) A phylogenetically-restricted essential cell cycle progression factor in the human pathogen Candida albicans. Nat Commun 13:4256. 10.1038/s41467-022-31980-3

Lattmann E, Deng T, Walser M, et al (2022) A DNA replication-independent function of pre-replication complex genes during cell invasion in C. elegans. PLoS Biol 20:e3001317. 10.1371/journal.pbio.3001317

Lau E, Tsuji T, Guo L, et al (2007) The role of pre-replicative complex (pre-RC) components in oncogenesis. The FASEB Journal 21:3786–3794. 10.1096/fj.07-8900rev

Legrand M, Jaitly P, Feri A, et al (2019) Candida albicans: An Emerging Yeast Model to Study Eukaryotic Genome Plasticity. Trends in Genetics 35:292–307. 10.1016/j.tig.2019.01.005

Lisby M, Mortensen UH, Rothstein R (2003) Colocalization of multiple DNA double-strand breaks at a single Rad52 repair centre. Nat Cell Biol 5:572–577. 10.1038/ncb997

Lisby M, Rothstein R, Mortensen UH (2001) Rad52 forms DNA repair and recombination centers during S phase. Proc Natl Acad Sci USA 98:8276–8282. 10.1073/pnas.121006298

Mangaonkar S, Nath S, Chatterji BP (2024) Microtubule dynamics in cancer metastasis: Harnessing the underappreciated potential for therapeutic interventions. Pharmacology & Therapeutics 263:108726. 10.1016/j.pharmthera.2024.108726

Meier SM, Steinmetz MO, Barral Y (2024) Microtubule specialization by +TIP networks: from mechanisms to functional implications. Trends in Biochemical Sciences 49:318–332. 10.1016/j.tibs.2024.01.005

Méndez J, Stillman B (2000) Chromatin Association of Human Origin Recognition Complex, Cdc6, and Minichromosome Maintenance Proteins during the Cell Cycle: Assembly of Prereplication Complexes in Late Mitosis. Molecular and Cellular Biology 20:8602–8612. 10.1128/MCB.20.22.8602-8612.2000

Mo L, Su B, Xu L, et al (2022) MCM7 supports the stemness of bladder cancer stem-like cells by enhancing autophagic flux. iScience 25:105029. 10.1016/j.isci.2022.105029

Montanaro L, Treré D, Derenzini M (2008) Nucleolus, Ribosomes, and Cancer. The American Journal of Pathology 173:301–310. 10.2353/ajpath.2008.070752

Nishitani H, Lygerou Z (2002) Control of DNA replication licensing in a cell cycle. Genes to Cells 7:523–534. 10.1046/j.1365-2443.2002.00544.x

Orsolic I, Jurada D, Pullen N, et al (2016) The relationship between the nucleolus and cancer: Current evidence and emerging paradigms. Seminars in Cancer Biology 37–38:36–50. 10.1016/j.semcancer.2015.12.004

Parker AL, Kavallaris M, McCarroll JA (2014) Microtubules and Their Role in Cellular Stress in Cancer. Front Oncol 4:. 10.3389/fonc.2014.00153

Parker MW, Bell M, Mir M, et al (2019) A new class of disordered elements controls DNA replication through initiator self-assembly. eLife 8:e48562. 10.7554/eLife.48562

Polisetty SD, Dutta S, Vadnala RN, et al (2025) Organization principles of dynamic three-dimensional genome architecture associated with centromere clustering states. Proc Natl Acad Sci USA 122:e2520310122. 10.1073/pnas.2520310122

Ren B, Yu G, Tseng GC, et al (2006) MCM7 amplification and overexpression are associated with prostate cancer progression. Oncogene 25:1090–1098. 10.1038/sj.onc.1209134

Reuss O, Vik Å, Kolter R, Morschhäuser J (2004) The SAT1 flipper, an optimized tool for gene disruption in Candida albicans. Gene 341:119–127. 10.1016/j.gene.2004.06.021

Reza MH, Dutta S, Goyal R, et al (2024) Expansion microscopy reveals characteristic ultrastructural features of pathogenic budding yeast species. Journal of Cell Science 137:jcs262046. 10.1242/jcs.262046

Sahoo B, Goyal R, Dutta S, et al (2025) *Candida albicans* : Insights into the Biology and Experimental Innovations of a Commonly Isolated Human Fungal Pathogen. ACS Infect Dis 11:1780–1815. 10.1021/acsinfecdis.5c00079

Sclafani RA, Holzen TM (2007) Cell Cycle Regulation of DNA Replication. Annu Rev Genet 41:237–280. 10.1146/annurev.genet.41.110306.130308

Shi Y-K, Yu YP, Zhu Z-H, et al (2008) MCM7 Interacts with Androgen Receptor. The American Journal of Pathology 173:1758–1767. 10.2353/ajpath.2008.080363

Shultz RW, Lee T-J, Allen GC, et al (2009) Dynamic Localization of the DNA Replication Proteins MCM5 and MCM7 in Plants. Plant Physiology 150:658–669. 10.1104/pp.109.136614

Sreekumar L, Kumari K, Guin K, et al (2021) Orc4 spatiotemporally stabilizes centromeric chromatin. Genome Res 31:607–621. 10.1101/gr.265900.120

Thakur J, Sanyal K (2012) A Coordinated Interdependent Protein Circuitry Stabilizes the Kinetochore Ensemble to Protect CENP-A in the Human Pathogenic Yeast Candida albicans. PLoS Genet 8:e1002661. 10.1371/journal.pgen.1002661

Tytell JD, Sorger PK (2006) Analysis of kinesin motor function at budding yeast kinetochores. The Journal of Cell Biology 172:861–874. 10.1083/jcb.200509101

Varma D, Chandrasekaran S, Sundin LJR, et al (2012) Recruitment of the human Cdt1 replication licensing protein by the loop domain of Hec1 is required for stable kinetochore–microtubule attachment. Nat Cell Biol 14:593–603. 10.1038/ncb2489

Varshney N, Sanyal K (2019) Nuclear migration in budding yeasts: position before division. Curr Genet 65:1341–1346. 10.1007/s00294-019-01000-x

Vassilev A, DePamphilis M (2017) Links between DNA Replication, Stem Cells and Cancer. Genes 8:45. 10.3390/genes8020045

Wang L, Liu X (2023) Pan-Cancer Multi-Omics Analysis of Minichromosome Maintenance Proteins (MCMs) Expression in Human Cancers. Front Biosci (Landmark Ed) 28:230. 10.31083/j.fbl2809230

Wargacki MM, Tay JC, Muller EG, et al (2010) Kip3, the yeast kinesin-8, is required for clustering of kinetochores at metaphase. Cell Cycle 9:2581–2588. 10.4161/cc.9.13.12076

Wattanathamsan O, Pongrakhananon V (2022) Emerging role of microtubule-associated proteins on cancer metastasis. Front Pharmacol 13:935493. 10.3389/fphar.2022.935493

Wu Y, Zhang Q, Lin Y, et al (2024) Replication licensing regulated by a short linear motif within an intrinsically disordered region of origin recognition complex. Nat Commun 15:8039. 10.1038/s41467-024-52408-0

Young MR, Tye BK (1997a) Mcm2 and Mcm3 are constitutive nuclear proteins that exhibit distinct isoforms and bind chromatin during specific cell cycle stages of Saccharomyces cerevisiae. MBoC 8:1587–1601. 10.1091/mbc.8.8.1587

Young MR, Tye BK (1997b) Mcm2 and Mcm3 are constitutive nuclear proteins that exhibit distinct isoforms and bind chromatin during specific cell cycle stages of Saccharomyces cerevisiae. Mol Biol Cell 8:1587–1601. 10.1091/mbc.8.8.1587

Yu S, Wang G, Shi Y, et al (2020) MCMs in Cancer: Prognostic Potential and Mechanisms. Analytical Cellular Pathology 2020:1–11. 10.1155/2020/3750294

Zasadzińska E, Huang J, Bailey AO, et al (2018) Inheritance of CENP-A Nucleosomes during DNA Replication Requires HJURP. Developmental Cell 47:348–362.e7. 10.1016/j.devcel.2018.09.003

Zhao Y, Wang J, Liang X, Wang C (2024) Clinical relevance of ORCs in predicting prognosis and immunotherapy outcomes: A pan-cancer analysis. Immunobiology 229:152783. 10.1016/j.imbio.2024.152783

Zheng D, Ye S, Wang X, et al (2017) Pre-RC Protein MCM7 depletion promotes mitotic exit by Inhibiting CDK1 activity. Sci Rep 7:2854. 10.1038/s41598-017-03148-3

