## Supplementary material for "Elevated Levels of Mcm7 Disrupt Microtubule and Nucleolar Homeostasis to Drive Genome Instability and Cell Death": All Supplementary data

**The file contains**

Supplementary Text

Supplementary Figures S1 to S5

Supplementary Tables S1 to S5

Supplementary references

### Supplementary Text

*Generation of overexpression strains*

All the overexpression plasmids used in this study (CIp10-$P_{TET}$: *EV, ORC3, ORC6, MCM4, MCM5, MCM7, ASE1*) were available from the library (Legrand et al. 2018; Jaitly et al. 2022). Plasmids were extracted using the boiling lysis method (Harwood 1996). Prior to transformation in the *C. albicans* strain containing the transactivator pNIMX, the plasmids were linearized with StuI. Correct transformants were PCR confirmed with PJ88/PJ89. All *C. albicans* transformations described hereafter were performed using the lithium acetate method (Walther and Wendland 2003).

*Generation of co-overexpression strains*

To co-overexpress Ase1 and Mcm7 in the same strain, the selection marker for CIp10-$P_{TET}$-*MCM7* was changed from Ura3 to HygB to overcome marker conflict. To achieve this, the *HYGB* was PCR-amplified using primers RG026/RG027 and incorporated into the NotI and XbaI sites of CIp10-$P_{TET}$-*MCM7*, which had been digested to release the *URA3* marker. Positive CIp10-$P_{TET}$-*MCM7-HYG* clones were verified by restriction digestion. The plasmid was then linearized with StuI and transformed into strains containing either the *EV* or CIp10-$P_{TET}$-*ASE1* to generate SRICa220-221 and SRICa225-226, respectively. To confirm the transformants, we performed multiplex PCR. PJ88/PJ89 confirmed the presence of *EV* or CIp10-$P_{TET}$-*ASE1* in the strain, as PJ89 binds to the *URA3* sequence. Integration of CIp10-$P_{TET}$-*MCM7-HYG* was confirmed using PJ88/RG027. Thus, the presence of two bands ensured the integration of both constructs in the same strain.

*Generation of mcm7 conditional mutant*

The first allele of *MCM7* was deleted using the pSFS2a plasmid vector backbone. An upstream sequence (US, sequence preceding the start codon) was amplified using primers SRI16/SRI17 and cloned into the KpnI/XhoI site of pSFS2a. Subsequently, a downstream sequence (DS, sequence after the stop codon) was amplified using primers SRI18/SRI19 and cloned into the SacI/SacII site of the US cloned vector, thereby generating pMcm7del. The cassette was confirmed by restriction digestion, and the resultant plasmid cassette was digested with KpnI and SacI for transformation into *C. albicans* cells. The heterozygous null transformants were confirmed by PCR using primers PJ3/SRI20.

The promoter of the second allele was replaced by the conditional $P_{MET3}$ promoter. Amplification of the 5' coding region of *MCM7,* starting from the start codon, was performed using primers SRI45/SRI46. The fragment was then cloned into the BamHI and PstI sites of pCaDis vector backbone to generate pMet3*MCM7*. Single-site integration in *C. albicans* genome was achieved by digesting the correct plasmid clone with BmgBI. PCR validation was performed using primers SRI34/SRI38.

*Tagging H4 and Rad52 with GFP*

We utilized the pFA-*TAP*-*ARG* plasmid backbone to integrate the *RP10* locus in the PvuI and SpeI sites to generate pFA-*TAP*-*ARG*_CIp10. This plasmid was further modified to replace the *TAP* epitope tag with GFP by digesting the backbone with NheI and SmaI, then inserting the GFP tag extracted from pGFP-HIS after digestion with SpeI and SmaI (NheI and SpeI have compatible ends). This resulted in pFA-GFP-*ARG*_CIp10. To tag H4 and Rad52, the promoter and *ORF* without the stop codon were amplified using primers SRI98/SRI99 (for H4) and SRI142/SRI143 (for Rad52). The amplified fragments were then cloned into the NotI and PacI sites of pFA-GFP-*ARG*_CIp10 to generate pH4-GFP-*ARG*_CIp10 and pRad52-GFP-*ARG*_CIp10, respectively. Correct clones were verified using restriction digestion. The plasmids were digested with StuI prior to transformation into *C. albicans* cells. Transformants were screened by fluorescence microscopy.

*Tagging of Mcm7 with fluorescent proteins*

The C-terminal 3' coding region of Mcm7 was amplified without the stop codon using the primers TEJ3_*MCM7*_SACII FP/TEJ4_*MCM7*_SPEI RP and cloned into the SacII and SpeI sites of pRFP-*ARG* to generate pMcm7-mCherry-*ARG*. The same fragment was extracted by digestion and then integrated into the SacII and SpeI sites of pGFP-*HIS* to generate pMcm7-GFP-*HIS*. Plasmids were verified by restriction digestion. The correct pMcm7-mCherry-*ARG* plasmid was linearized with XhoI for transformation into *C. albicans* to generate SRICa075-077 and SRICa082. Due to the occurrence of two XhoI sites in the pMcm7-GFP-HIS, the plasmid was partially digested with XhoI, followed by transformation to generate SRICa244. The correct *C. albicans* transformants were screened by fluorescence microscopy and western blotting.

To generate CIp10-$P_{TET}$-*MCM7*-GFP-*URA3*, we digested the pCIp10-$P_{TET}$-*GTW* with EcoRV and cloned GFP using the primers SRI82/SRI83. The resultant plasmid was then digested with PacI and PciI to insert PTETMCM7 (the *MCM7 ORF* along with the $P_{TET}$ promoter), which was amplified using SRI148/SRI149. The plasmid was then linearized with StuI for transforming the *C. albicans* strains, and the correct transformants were PCR confirmed using PJ88/PJ89.

*Tagging of Nop1 with mCherry*

The C-terminus of Nop1 was tagged with mCherry by amplification of the 3' coding region of Nop1 without the stop codon using primers SRI90/SRI91. The fragment was cloned into the SacII and SpeI sites of pRFP-ARG4 to generate pNop1-mCherry-ARG. Post confirmation with restriction digestion, the plasmid was partially digested with MfeI and transformed into *C. albicans* cells to generate SRICa243. The transformants were screened by fluorescence microscopy.

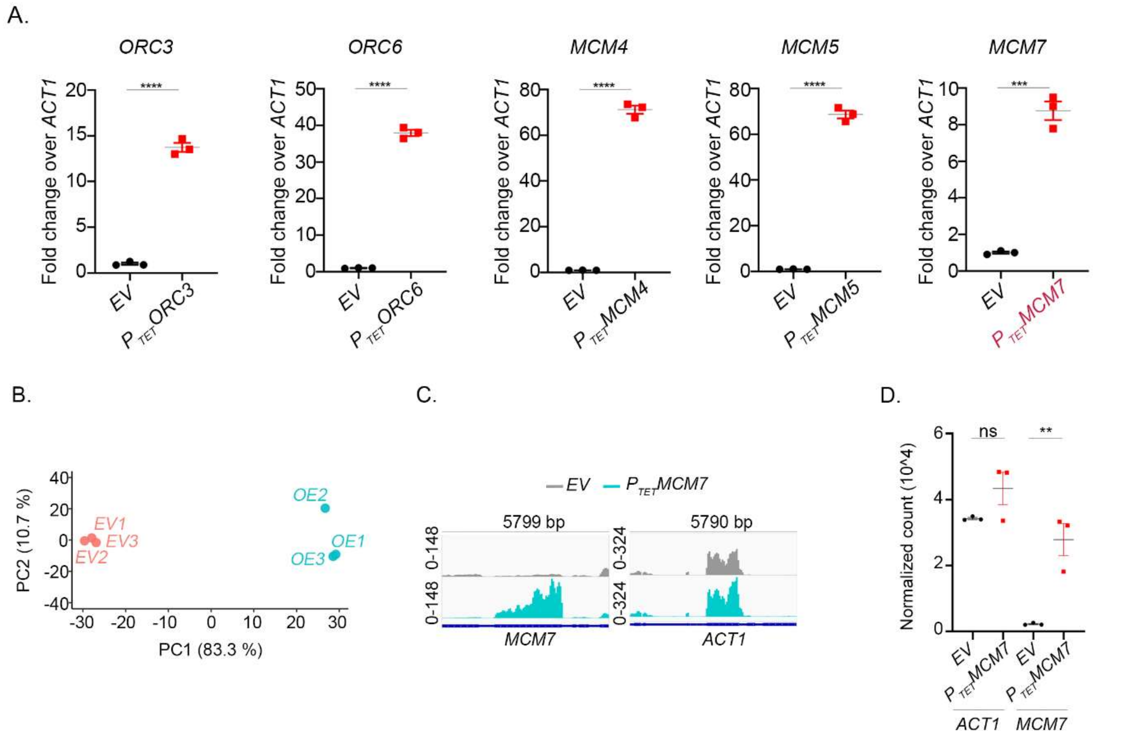


**Figure S1. Elevated Mcm7 levels alter the transcriptional landscape in *C. albicans*.** A) Real-time PCR quantification of pre-RC transcripts in *EV* (CaPJ160) and respective overexpression strains ($P_{TET}$*ORC3*- SRICa007; $P_{TET}$*ORC6*- SRICa012; $P_{TET}$*MCM4*- SRICa016; $P_{TET}$*MCM5*- SRICa020; $P_{TET}$*MCM7*- CaPJ165), all treated with Dox for 8 h. Fold change relative to the housekeeping gene actin (*ACT1*) has been plotted. $N = 3$, unpaired *t*-test, *P*-value: (****) < 0.0001, (***) <0.001. B) Principal component analysis (PCA) plot highlighting variation amongst and within groups analyzed in the RNA-seq experiment. *EV* (1-3) and *OE* (1-3) represent the three biological transformants for the empty vector control (CaPJ170) and Mcm7 overexpression (CaPJ165), respectively. C) Snapshot of *MCM7* and *ACT1* transcript levels as viewed in the integrative genomics viewer (IGV) for a single biological transformant. D) Plot representing normalized transcript counts from the analysed RNA-seq data. $N = 3$, two-way ANOVA with Tukey's multiple comparisons test, *P*-value: (**) < 0.01, (ns) >0.05. All error bars indicate the standard error of mean (SEM).

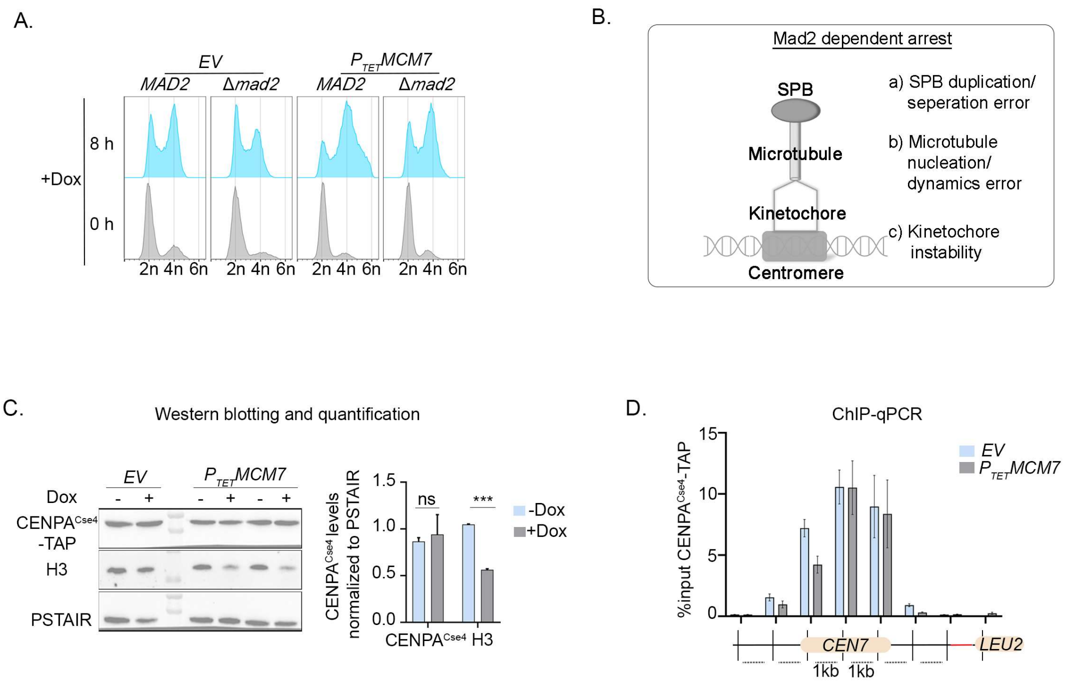


**Figure S2. Excess Mcm7 impacts histone H3 levels but not CENPA$^{Cse4}$ stability.** A) FACS profile at 0 h and 8 h post-induction with Dox to assay for DNA content upon Mcm7 overexpression, with (SRICa064) and without active SAC (SRICa131). Cyclic population in *EV*, both with (CaPJ170) and without (SRICa246) active SAC, serves as a control. B) Schematic highlighting key biological processes, dysregulation of which often activates the Mad2-dependent SAC, an event triggered upon Mcm7 overexpression. C) (*left*) Western blot and (*right*) quantification to measure cellular levels of CENPA$^{Cse4}$-TAP and H3 in Mcm7 overexpression condition (SRICa064), $N = 2$, paired *t*-test, *P*-value: (**) < 0.01, (ns) > 0.05. PSTAIR levels are used as a loading control. D) ChIP-qPCR values of CENPA$^{Cse4}$-TAP probed at centromere 7 (*CEN7*) and a non-centromeric region control (*LEU2*) on chr 7. At *CEN7*, primer pairs were designed at approximately 1-kb intervals to scan ~ 6 kb of the centromeric region (as indicated by the vertical lines at the bottom). Primer pairs: nCEN (-1,0), (1,2), (3,4), (5,6), (7,8) from left to right. $N = 3$; two-way ANOVA; Sidak's multiple-comparison test applied between *EV* and $P_{TET}$*MCM7* strains for each primer pair; non-significant for each comparison. All error bars indicate SEM.

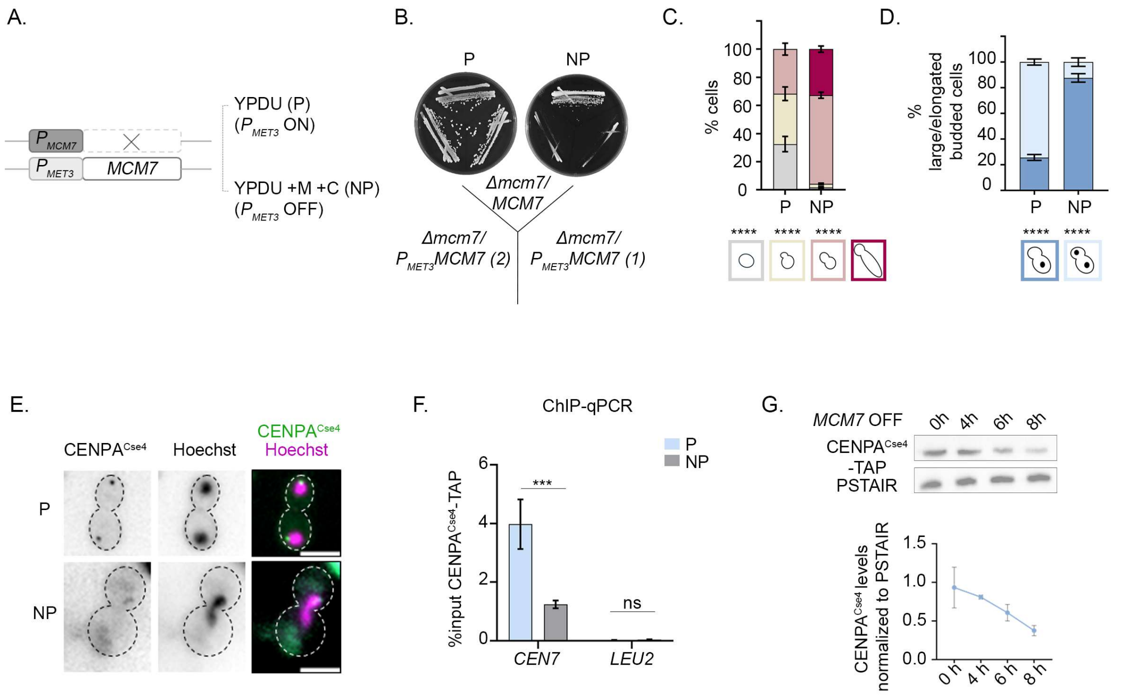


**Figure S3. Mcm7 is essential for cell viability and CENPA$^{Cse4}$ stability.** A) Schematic representation of the conditional depletion strategy of Mcm7. The addition of methionine (+M) and cysteine (+C) to the media represses the +M +C-sensitive promoter ($P_{MET3}$), effectively depleting protein levels with each round of cell division. B) Streaking Mcm7 depletion strains (SRICa041 and SRICa045) on YPDU or permissive media (P) and YPDU +M +C or non-permissive media (NP) to assess cell lethality upon depletion of Mcm7 (on NP). *Δmcm7/MCM7* acts as the heterozygous null control (SRICa033). Plate photographed after 48 h of incubation at 30 °C. C) and D) Quantification for the distribution of different cell cycle stages and nuclear segregation pattern, respectively, in the presence and absence of Mcm7. $N = 3$, $n \geq 100$ for each biological replicate. 2-way ANOVA with Sidak's multiple comparisons test, (****) P-value<0.0001, (ns) P-value>0.05. E) Fluorescence microscopy images of CENPA$^{Cse4}$-GFP in large-bud cells in the presence or absence of Mcm7. The nucleus is stained with Hoechst. Scale bar: 5 µm. F) ChIP-qPCR values of CENPA$^{Cse4}$-TAP probed at centromere 7 (*CEN7*) and a non-centromeric region control (*LEU2*) on chr 7 in Mcm7 ON and OFF conditions. $N = 2$, 2-way ANOVA with Sidak's multiple comparisons test, (***) P-value<0.001, (ns) > 0.05. G) (*top*) Western blot and (*bottom*) quantification to measure cellular levels of CENPA$^{Cse4}$-TAP at different time points of Mcm7 depletion. All error bars indicate SEM.

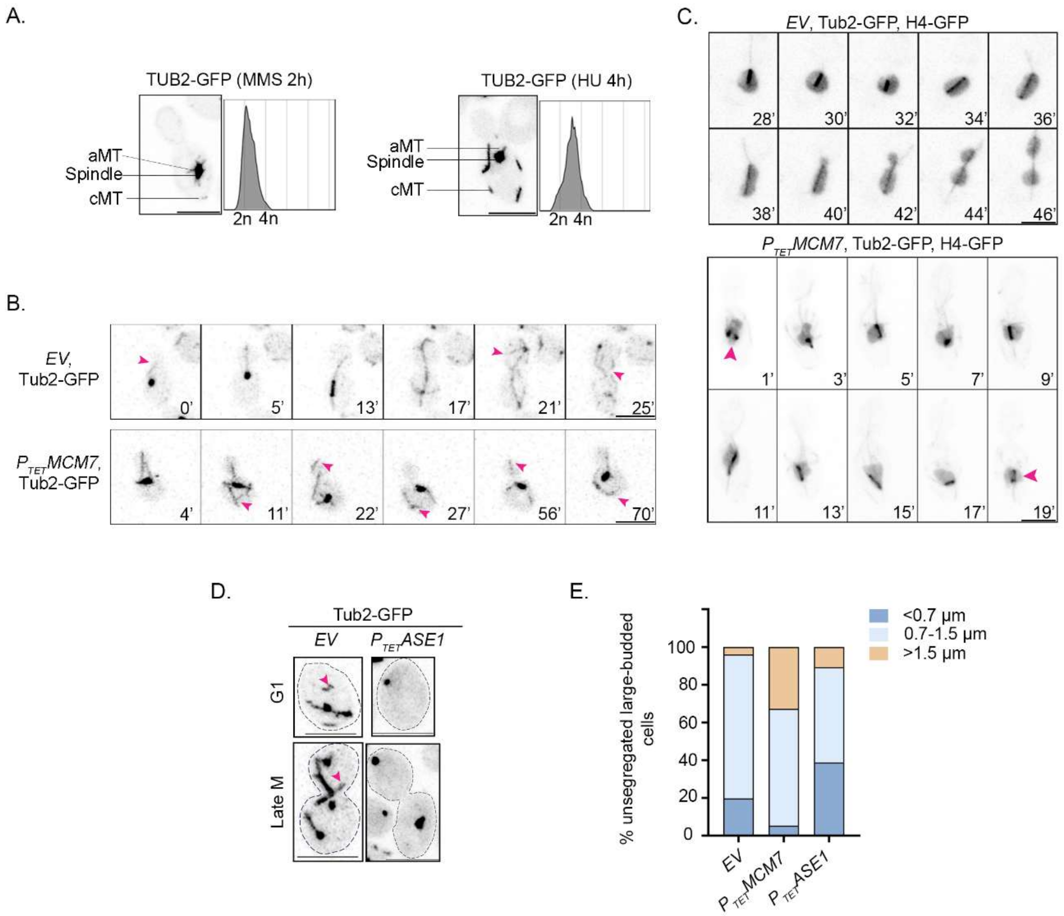


**Figure S4. Mcm7 overexpression perturbs microtubule organization and elicits defects contrasting with Ase1 overexpression.** A) MT landscape as observed by fluorescence microscopy of Tub2-GFP in $P_{TET}$*MCM7* cells (SRICa157) treated with MMS and HU for the indicated time. FACS profile depicting cell cycle arrest in G1/S phase due to replication block. B) Live-cell microscopy montages of Tub2-GFP to chase MT dynamics in (*top*) *EV* (CaRG080) and (*below*) $P_{TET}$*MCM7* (SRICa157) treated with Dox. Pink arrowheads indicate cMTs. The bottom corner represents time in min. Scale bar: 5 µm. C) Live-cell microscopy montages of H4-GFP representing the nucleus (lighter diffuse signals) and Tub2-GFP representing the MTs (dark rod-shaped signal marks the spindle) in *EV* (SRICa208) and $P_{TET}$*MCM7* cells (SRICa211) treated with Dox. Pink arrowheads indicate instances of spindle buckling. The lower corner indicates time in min. Scale bar: 5 µm. D) Fluorescence microscopy of Tub2-GFP, highlighting the status of cMTs (pink arrowheads) in *EV* (CaRG080) and Ase1 overexpression ($P_{TET}$*ASE1*- SRICa236). Scale bar: 5 µm. E) Distribution of percentage large-budded cells, unsegregated nucleus with the indicated spindle length in *EV* (CaRG080), $P_{TET}$*MCM7* (SRICa211), and $P_{TET}$*ASE1* (SRICa236).

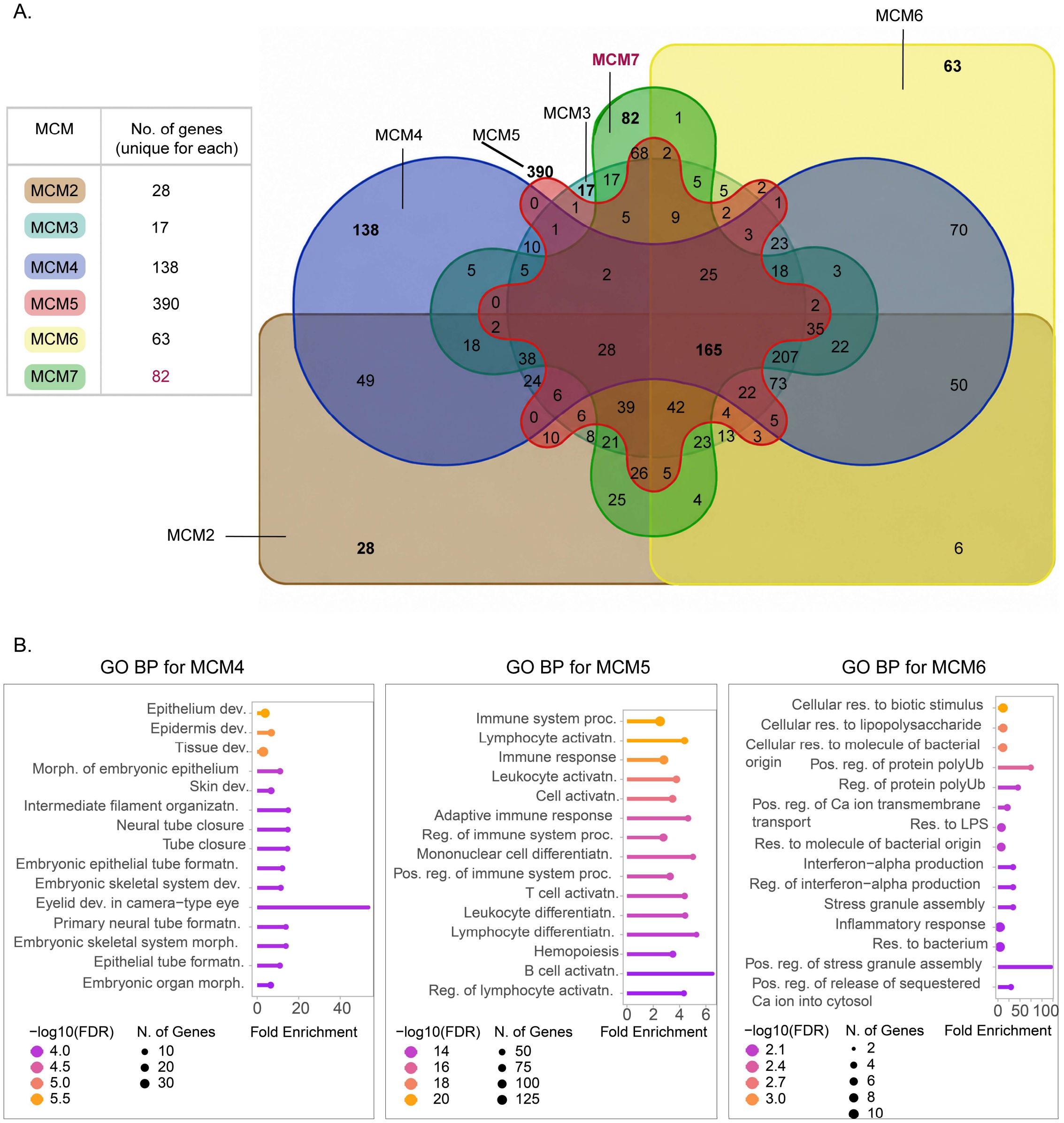


Abbreviations: (GO) Gene Ontology, (BP) Biological Processes, (dev.) development, (morph.) morphogenesis, (organizatn.) organization, (formatn.) formation, (proc.) processes, (activatn.) activation, (reg.) regulation, (pos.) positive, (differentiatn.) differentiation, (res.) response.

**Figure S5. Distinct biological processes associated with dysregulation of individual MCM subunits in cancer.** A) Multi-set overlap analysis of gene sets whose expression is positively correlated with the expression of MCM subunits in cancer, and listing the number of genes uniquely perturbed for each MCM subunit. B) Gene ontology (GO) terms, Biological Processes (BP), for the gene sets uniquely associated with each MCM subunit. GO-BP for MCM2 and MCM3 could not be obtained, possibly due to the very low number of genes uniquely associated with them (28 for MCM2 and 17 for MCM3). GO BP for MCM7 is shown in Fig. 5G. GO analysis was conducted in ShinyGO 0.85.1.

**Supplementary Table S1.** List of strains used in this study

| Name | Description | Genotype | Reference |
|---|---|---|---|
| SN148 | Wild type | *Δura3::imm434/Δura3::imm434, Δhis1::hisG/Δhis1::hisG, Δarg4::hisG/Δarg4::hisG, Δleu2::hisG/Δleu2::hisG* | (Noble and Johnson 2005) |
| YJB8675 | *CSE4-GFP* | *Δura3::imm434/Δura3::imm434, Δhis1::hisG/Δhis1::hisG, Δarg4::hisG/Δarg4::hisG, CSE4-GFPCSE4/CSE4* | (Joglekar et al. 2008) |
| J110 | *mad2* deletion in SN148 | SN148 *mad2::ARG4/mad2::LEU2* | (Thakur and Sanyal 2011) |
| CaPJ158 | transactivator in *CSE4-GFP* | YJB8675 *ADH1/adh1::PTDH3-cartTA-SAT1* | (Jaitly et al. 2022) |
| CaPJ159 | transactivator in *CSE4-GFP, TUB4-mCherry* | YJB8675 *ADH1/adh1::*$P_{TDH3}$*-cartTA-SAT1*, *TUB4/TUB4-mCherry-ARG4* | (Jaitly et al. 2022) |
| CaPJ160 | *EV* in *CSE4-GFP, TUB4-mCherry* | CaPJ159 *RPS1/RPS1::*$P_{TET}$*-GtwB-URA3* | (Jaitly et al. 2022) |
| CaPJ165 | $CSA4^{MCM7}$ overexpression in *CSE4-GFP, TUB4-mCherry* | CaPJ159 *RPS1/RPS1::*$P_{TET}$*-MCM7-URA3* | (Jaitly et al. 2022) |
| CaPJ169 | SNX1or transactivator in SN148 | SN148 *ADH1/adh1::*$P_{TDH3}$*-cartTA-SAT1* | (Jaitly et al. 2022) |
| CaPJ170 | *EV* in SNX1 | CaPJ169 *RPS1/RPS1::*$P_{TET}$*-GtwB-URA3* | (Jaitly et al. 2022) |
| CaPJ196 | transactivator in *mad2* deletion | J110 *ADH1/adh1::*$P_{TDH3}$*-cartTA-SAT1* | (Jaitly et al. 2022) |
| CaRG080 | *EV* in *TUB2-GFP, TUB4-mCherry* | CaPJ170 *TUB2/TUB2-GFP-HIS1, TUB4/TUB4-mCherry-ARG4* | (Reza et al. 2024) |
| CaRG090 | *TUB2-GFP* in SNX1 | CaPJ169 *TUB2/TUB2-GFP-HIS1* | This study |
| CaRG095 | *TUB2-GFP, TUB4-mCherry* in SNX1 | CaRG090 *TUB4/TUB4-mCherry-ARG4* | This study |
| SRICa007 | *ORC3* overexpression in *CSE4-GFP* | CaPJ158 *RPS1/RPS1::*$P_{TET}$*-ORC3-URA3* | This study |

| | | | |
|---|---|---|---|
| SRICa012 | *ORC6* overexpression in *CSE4-GFP* | CaPJ158 *RPS1/RPS1::*$P_{TET}$*-ORC6-URA3* | This study |
| SRICa016 | *MCM4* overexpression in *CSE4-GFP* | CaPJ158 *RPS1/RPS1::*$P_{TET}$*-MCM4-URA3* | This study |
| SRICa020 | *MCM5* overexpression in *CSE4-GFP* | CaPJ158 *RPS1/RPS1::*$P_{TET}$*-MCM5-URA3* | This study |
| SRICa051 | *CSE4-TAP* in SNX1 | CaPJ169 *CSE4/CSE4-TAP-LEU2* | This study |
| SRICa064 | *MCM7* overexpression in *CSE4-TAP* | SRICa051 *RPS1::*$P_{TET}$*-MCM7-URA3/RPS1* | This study |
| SRICa175 | *H4-GFP* in *EV* control | CaPJ170 *RPS1::*$P_{TET}$*-GtwB-URA3/RPS1::H4-GFP-ARG4* | This study |
| SRICa178 | *H4-GFP* in *MCM7* overexpression | SRICa064 *RPS1::*$P_{TET}$*-MCM7-URA3/RPS1::H4-GFP-ARG4* | This study |
| SRICa234 | *RAD52-GFP* in *MCM7* overexpression | SRICa064 *RPS1::*$P_{TET}$*-MCM7-URA3/RPS1::RAD52-GFP-ARG4* | This study |
| SRICa265 | *RAD52-GFP* in *EV* control | CaPJ170 *RPS1::*$P_{TET}$*-GtwB-URA3/RPS1::RAD52-GFP-ARG4* | This study |
| SRICa246 | *EV* in *mad2* deletion | CaPJ196 *RPS1/RPS1::*$P_{TET}$*-GtwB-URA3* | This study |
| SRICa131 | *MCM7* overexpression in *mad2* deletion | CaPJ196 *RPS1/RPS1::*$P_{TET}$*-MCM7-URA3* | This study |
| SJCa001 | *CSE4-TAP* in SN148 | SN148 *CSE4/CSE4-TAP-LEU2* | This study |
| SRICa033 | *MCM7* heterozygous null in *CSE4-GFP* | YJB8675 *mcm7::FRT/MCM7* | This study |
| SRICa035 | *MCM7* heterozygous null in *CSE4-TAP* | SJCa001 *mcm7::FRT/MCM7* | This study |
| SRICa041 | *mcm7* conditional mutant in *CSE4-GFP* | SRICa033 *mcm7::FRT/*$P_{MET3}$*MCM7-URA3* | This study |
| SRICa045 | *mcm7* conditional mutant in *CSE4-TAP* | SRICa035 *mcm7::FRT/*$P_{MET3}$*MCM7-URA3* | This study |
| SRICa157 | *MCM7* overexpression in *TUB2-GFP, TUB4-mCherry* | CaRG095 *RPS1/RPS1::*$P_{TET}$*-MCM7-URA3* | This study |
| SRICa208 | *TUB2-GFP* in *EV*, *H4-GFP* | SRICa175 *TUB2/TUB2-GFP-HIS1* | This study |

| | | | |
|---|---|---|---|
| SRICa211 | *TUB2-GFP* in *MCM7* overexpression, *H4-GFP* | SRICa178 *TUB2/TUB2-GFP-HIS1* | This study |
| SRICa236 | *ASE1* overexpression in *TUB2-GFP, TUB4-mCherry* | CaRG095 *RPS1/RPS1::*$P_{TET}$*-ASE1-URA3* | This study |
| SRICa245 | *ASE1* overexpression in *CSE4-GFP* | CaPJ159 *RPS1/RPS1::*$P_{TET}$*-ASE1-URA3* | This study |
| SRICa220 | *EV* in *MCM7* overexpression | CaRG080 *RPS1::*$P_{TET}$*-GtwB-URA3/RPS1::*$P_{TET}$*-MCM7-HYG* | This study |
| SRICa225 | *MCM7-ASE1* co-overexpression | SRICa245 *RPS1::*$P_{TET}$*-ASE1-URA3/RPS1::*$P_{TET}$*-MCM7-HYG* | This study |
| SRICa239 | *MCM7-GFP* overexpression | SRICa051 *RPS1/RPS1::*$P_{TET}$*-MCM7-GFP-URA3* | This study |
| SRICa244 | *MCM7-GFP* native expression | SN148 *MCM7/MCM7-GFP-HIS1* | This study |
| SRICa243 | *NOP1-mCherry* in *MCM7-GFP* overexpression | SRICa239 *NOP1/NOP1-mCherry-ARG4* | This study |
| SRICa075 | *MCM7-mCherry* native expression | SRICa035 *mcm7::FRT/MCM7-mCherry-ARG4* | This study |
| SRICa082 | *MCM7-mCherry* overexpression | SRICa064 *RPS1/RPS1::*$P_{TET}$*-MCM7-mCherry-ARG4-URA3* | This study |

**Supplementary Table S2.** List of plasmids used in this study

| **Name** | **Description** | **Reference** |
|---|---|---|
| CIp10-$P_{TET}$-GTW derivatives | Overexpression plasmid collection | (Legrand et al. 2018) |
| pNIMX | Plasmid harbouring $P_{TET}$ transactivator | (Chauvel et al. 2012) |
| pCse4-TAP-LEU | TAP-tagging plasmid for Cse4 | (Jaitly et al. 2022) |
| pGFP-HIS | GFP-tagging plasmid | (Chatterjee et al. 2016) |
| pRFP-ARG | mCherry-tagging plasmid | (Varshney and Sanyal 2019) |
| pTub2-GFP-HIS | GFP-tagging plasmid for Tub2 | (Reza et al. 2024) |
| pFA-TAP-ARG | TAP-tagging plasmid | (Lavoie et al. 2008) |
| pSFS2a | Recyclable *SAT1*-flipper cassette | (Reuss et al. 2004) |
| pCaDis | Plasmid for promoter replacement with $P_{MET3}$ | (Care et al. 1999) |
| pFA-TAP-ARG_CIp10 | TAP-tagging plasmid with *RP10* locus | This study |
| pFA-GFP-ARG_CIp10 | GFP-tagging plasmid with *RP10* locus | This study |
| pH4-GFP-ARG_CIp10 | GFP-tagging plasmid for H4 | This study |
| pRad52-GFP-ARG_CIp10 | GFP-tagging plasmid for Rad52 | This study |
| pMcm7del | Deletion cassette for *MCM7* | This study |
| pMet3Mcm7 | Plasmids for promoter replacement of *MCM7* with *MET3* | This study |
| CIp10-$P_{TET}$-MCM7-HYG | Overexpression plasmid for Mcm7 with HygB | This study |
| pMcm7-mCherry-ARG | mCherry-tagging plasmid for Mcm7 | This study |
| pMcm7-GFP-HIS | GFP-tagging plasmid for Mcm7 | This study |

| | | |
|---|---|---|
| CIp10-$P_{TET}$-MCM7-GFP-URA | Overexpression plasmid for Mcm7-GFP | This study |
| pNop1-mCherry-ARG | mCherry-tagging plasmid for Nop1 | This study |

**Supplementary Table S3.** List of primers used in this study

| Name | Sequence | Description |
|---|---|---|
| PJ88 | ATACTACTGAAAATTTCCTGACTTTC | Confirmation of overexpression plasmid integration |
| PJ89 | ATTACTATTTACAATCAAAGGTGGTC | |
| RG026 | ATAGCGGCCGCGTATAGTGCTTGCTGTTCGATATTG | Amplification of HygB |
| RG027 | GCGTCTAGAATTTTATGATGGAATGAATGGGATG | |
| SRI16 | CATGGTACCGGTCCTGAATCTCTTCGTAAG | Amplification of *MCM7* US |
| SRI17 | CGTCTCGAGCCTCTTTCCAATGTGGTGG | |
| SRI18 | GAACCGCGGGTTGAAGCAGGAGGATTCT | Amplification of *MCM7* DS |
| SRI19 | CTGGAGCTCCTTTTTGGGAAATTTAGACGTAC | |
| PJ3 | CTATTCTCTAGAAAGTATAGGAACTTC | Confirmation of *MCM7* first allele deletion |
| SRI20 | ACGATACTGATGAAGAGGATTG | |
| SRI45 | GTACTGGATCCATGTCAACCACTACGGCAGCTG | Cloning of *MCM7* under *MET3* |
| SRI46 | GGCTACTGCAGTAACATCCAACACATCATCACGGTAAC | |
| SRI34 | GCTAGCTGCAGAATTGTCTATTCCAAGCCTGTG | Confirmation of *MCM7* under *MET3* |
| SRI38 | CCAGTGTATGGACTTGGCATG | |
| SRI98 | GTCGCAGCGGCCGCGAGAGAGAGACCAAAAAGTGGC | GFP tagging of H4 |
| SRI99 | GCTGCTTTAATTAAACCACCGAAACCATACAAGGTTC | |
| SRI142 | TGTCGAGCGGCCGCCTCATATTCAAAATTCTAGGTGG | GFP tagging of Rad52 |
| SRI143 | GTGCATTTAATTAATTGGTTAACAGTCGTATTAGC | |
| TEJ3_MCM7_SACII FP | ATTACCGCGGCCAAGATTGTCTCCACATG | C-term tagging of Mcm7 with mCherry |
| TEJ4_MCM7_SPEI RP | ATGCACTAGTCAATATTAAAAGATTCTCTCCATCATC | |
| SRI82 | CCTGATGATATCATGAATAAACTTCCCAAAGGAT | |

| | | |
|---|---|---|
| SRI83 | CCTGCAGATATCCTGTGCAATAACTTTCTGTCC | Cloning of GFP in overexpression vector |
| SRI148 | GTCGTAACATGTTCTTTCCTGCGTTATC | Cloning of *MCM7* in overexpression-GFP plasmid |
| SRI149 | CGTGCATTAATTAACAATATTAAAAGATTCTCTCCATCATC | |
| SRI90 | CATACTCCGCGGCTCTCATCGTCCAGGTAGAG | C-term tagging of Nop1 with mCherry |
| SRI91 | GTACTGACTAGTTTTCTTTATTCCGCTTCTCATG | |
| Primers for real-time PCR | | |
| SRI114 | CAATTTATACGACTTTTATCAGG | For *ORC3* transcript |
| SRI115 | GGACAAACCACGCATAAG | |
| SRI116 | GAAATTCTTTTGAAAGAAAAGCG | For *ORC6* transcript |
| SRI117 | CATTTCTCTAAAGCCTCTTTACG | |
| SRI118 | CAAACTGGTACTACGGCAC | For *MCM4* transcript |
| SRI119 | CCACTCTAAATGAACTACGTTC | |
| SRI120 | CTTGGAAGCTCGAATGATGTTAC | For *MCM5* transcript |
| SRI121 | CTTTACGCAATGTCTTATAAGC | |
| SRI122 | GCCATTAGATTAATTGAAGTGAG | For *MCM7* transcript |
| SRI123 | TTGATTTGATCTAATGCAACTCG | |
| SRI124 | CAAACCATCCAAGATAGTTCTTC | For *CDT1* transcript |
| SRI125 | TCATATCAAGTTTTGACGGTAG | |
| SRI126 | TTGGCTCCATCTTCTATGAAAG | For *ACT1* transcript |
| SRI127 | CATTTGTTGGAAAGTAGACAATG | |
| nCEN(-1) | GACGTTTCTCAAATCATATAACATTC | *CEN7* ChIP qPCR primers |
| nCEN(0) | ACCACTGGAGCTTTCCAGTAG | |
| nCEN(1) | CACCTCTGCACTAATCTACA | |

| | | |
|---|---|---|
| nCEN(2) | TGTTGAGATGTCTTATTGAT | |
| nCEN(3) | GCATACCTGACACTGTCGTT | |
| nCEN(4) | AACGGTGCTACGTTTTTTTA | |
| nCEN(5) | TCAATTATCGCTTGATAGCG | |
| nCEN(6) | CTATCATCATGCCAGCCTAG | |
| nCEN(7) | CAGAGCAATGGCCCTTGTG | |
| nCEN(8) | GCTTACTCAATCTACTTTGTCTTCCA | |
| nLEU2(1) | GTACCGAAATTGTCAATGAAG | *LEU2* ChIP qPCR primers |
| nLEU2(2) | GTGGTGTTTGAAATCAAATTG | |

**Supplementary Table S4.** List of human cell lines

| Name | Source | Identifier |
|---|---|---|
| NCI-H1299 | ATCC | ATCC® CRL-5803™ |
| HepG2 | ATCC | ATCC® HB-8065™ |

**Supplementary Table S5.** List of chemicals and antibodies

| Name | Source | Identifier |
|---|---|---|
| Antibodies | | |
| Polyclonal anti-Protein A | Sigma | Cat. no. P3775 |
| Monoclonal anti-PSTAIR | Sigma | Cat. no. P7962 |
| Monoclonal anti-RFP | ChromoTek | Cat. no. 5F8-100 |
| Polyclonal anti-H3 | Abcam | Cat. no. AB1791 |
| Polyclonal anti-MCM7 | ABclonal | Cat. no. A1138 |
| Monoclonal anti-NPM1 | In-house generated | (Senapati et al. 2022) |
| Monoclonal anti-β-Actin | ABclonal | Cat. no. AC026 |
| Goat anti-rabbit IgG-HRP | Abcam | Cat. no. AB97051 |
| Goat anti-mouse IgG-HRP | Abcam | Cat. no. AB97023 |
| Rabbit anti-rat IgG-HRP | Genei | Cat. no. HPO14 |
| Alexa Fluor 488 goat anti-rabbit IgG (H+L) | Invitrogen | Cat. no. A11001 |
| Alexa Fluor 568 goat anti-mouse IgG (H+L) | Invitrogen | Cat. no. A11011 |
| Reagents | | |
| Doxycycline hyclate | Sigma | Cat. no. D9891 |
| Hoechst 33342 | Sigma | Cat. no. B2261 |
| DAPI | Merck | Cat. no. D9542 |
| Hydroxyurea | HiMedia | Cat no. 6487 |
| Methyl methanesulfonate | Merck | Cat no. 129925 |
| Protein A-Sepharose beads | Merck | Cat no. P9424 |
| Zymolyase-20T | MP Biomedicals | Cat no. 32092 |
| TRIzol Reagent | Invitrogen | Cat no. 15596026 |
| Nourseothricin | Jena bioscience | Cat no. AB-102XL |

### References

Care RS, Trevethick J, Binley KM, Sudbery PE (1999) The *MET3* promoter: a new tool for *Candida albicans* molecular genetics. Molecular Microbiology 34:792–798. https://doi.org/10.1046/j.1365-2958.1999.01641.x

Chatterjee G, Sankaranarayanan SR, Guin K, et al (2016) Repeat-Associated Fission Yeast-Like Regional Centromeres in the Ascomycetous Budding Yeast Candida tropicalis. PLoS Genet 12:e1005839. https://doi.org/10.1371/journal.pgen.1005839

Chauvel M, Nesseir A, Cabral V, et al (2012) A Versatile Overexpression Strategy in the Pathogenic Yeast Candida albicans: Identification of Regulators of Morphogenesis and Fitness. PLoS ONE 7:e45912. https://doi.org/10.1371/journal.pone.0045912

Harwood AJ (1996) The Rapid Boiling Method for Small-Scale Preparation of Plasmid DNA. In: Basic DNA and RNA Protocols. Humana Press, New Jersey, pp 265–268

Jaitly P, Legrand M, Das A, et al (2022) A phylogenetically-restricted essential cell cycle progression factor in the human pathogen Candida albicans. Nat Commun 13:4256. https://doi.org/10.1038/s41467-022-31980-3

Joglekar AP, Bouck D, Finley K, et al (2008) Molecular architecture of the kinetochore-microtubule attachment site is conserved between point and regional centromeres. The Journal of Cell Biology 181:587–594. https://doi.org/10.1083/jcb.200803027

Lavoie H, Sellam A, Askew C, et al (2008) A toolbox for epitope-tagging and genome-wide location analysis in Candida albicans. BMC Genomics 9:578. https://doi.org/10.1186/1471-2164-9-578

Legrand M, Bachellier-Bassi S, Lee KK, et al (2018) Generating genomic platforms to study Candida albicans pathogenesis. Nucleic Acids Research. https://doi.org/10.1093/nar/gky747

Noble SM, Johnson AD (2005) Strains and Strategies for Large-Scale Gene Deletion Studies of the Diploid Human Fungal Pathogen *Candida albicans*. Eukaryot Cell 4:298–309. https://doi.org/10.1128/EC.4.2.298-309.2005

Reuss O, Vik Å, Kolter R, Morschhäuser J (2004) The SAT1 flipper, an optimized tool for gene disruption in Candida albicans. Gene 341:119–127. https://doi.org/10.1016/j.gene.2004.06.021

Reza MH, Dutta S, Goyal R, et al (2024) Expansion microscopy reveals characteristic ultrastructural features of pathogenic budding yeast species. Journal of Cell Science 137:jcs262046. https://doi.org/10.1242/jcs.262046

Senapati P, Bhattacharya A, Das S, et al (2022) Histone Chaperone Nucleophosmin Regulates Transcription of Key Genes Involved in Oral Tumorigenesis. Mol Cell Biol 42:e00669-20. https://doi.org/10.1128/mcb.00669-20

Thakur J, Sanyal K (2011) The Essentiality of the Fungus-Specific Dam1 Complex Is Correlated with a One-Kinetochore-One-Microtubule Interaction Present throughout the Cell Cycle, Independent of the Nature of a Centromere. Eukaryot Cell 10:1295–1305. https://doi.org/10.1128/EC.05093-11

Varshney N, Sanyal K (2019) Aurora kinase Ipl1 facilitates bilobed distribution of clustered kinetochores to ensure error-free chromosome segregation in *Candida albicans*. Molecular Microbiology 112:569–587. https://doi.org/10.1111/mmi.14275

Walther A, Wendland J (2003) An improved transformation protocol for the human fungal pathogen Candida albicans. Curr Genet 42:339–343. https://doi.org/10.1007/s00294-002-0349-0
